# Study of Protein-Protein Interactions in Septin Assembly: Multiple amphipathic helix domains cooperate in binding to the lipid membrane

**DOI:** 10.1101/2025.08.08.669402

**Authors:** S. Mahsa Mofidi, Abhilash Sahoo, Christopher J. Edelmaier, Stephen J. Klawa, Ronit Freeman, Amy Gladfelter, M. Gregory Forest, Ehssan Nazockdast, Sonya M. Hanson

## Abstract

Septins are a conserved family of cytoskeletal proteins known for sensing micron-scale membrane curvature via amphipathic helix (AH) domains. While cooperative interactions in septin assembly have been suggested, the molecular mechanisms governing membrane binding and assembly remain unclear. Building on prior findings, we use all-atom molecular dynamics simulations to examine how single and paired extended AH domains, derived from Cdc12, interact with lipid bilayers. A single membrane-bound AH adopts a curved conformation. In solution, a second AH peptide preferentially interacts with the bound peptide through conserved salt bridges, favoring an antiparallel arrangement. Simulations of covalently linked AH tandems confirm this configuration. Dual membrane-bound peptides induce lipid packing defects, reduce tail order, and exhibit slight membrane displacement, suggesting curved membranes may better accommodate multiple AH domains. Our findings advance the mechanistic understanding of septin-membrane interactions and highlight the role of cooperative AH domain binding in stabilizing higher-order structures.

## INTRODUCTION

Septins are a conserved family of GTP-binding proteins that play essential roles in various cellular processes, including cell division, membrane remodeling, and cytoskeletal organization (*1*, *2*). A hallmark of septin function is their ability to sense and respond to membrane curvature, a property critical for maintaining cellular architecture and enabling dynamic shape changes (*3*). Septins polymerize into filaments and higher-order assemblies (*4*) that can scaffold and remodel cellular membranes (*5*). In *Saccharomyces cerevisiae*, septins form hetero-oligomeric complexes that polymerize along the membrane (*6*, *7*). Previous studies revealed that amphipathic helices (AHs) at the C-terminal end of the Cdc12 subunit shallowly insert into the lipid bilayer and facilitate membrane binding (*8*). Experimental studies have shown that septins’ AH domains bind to membranes and sense micron-scale curvature (*9*, *10*). A kinetic model showed septins undergo a multi-step assembly process, and curvature preference is modulated by protein concentration and membrane geometry (*11*). Although kinetic models suggest cooperative septin-membrane interactions, the molecular mechanisms underlying this cooperativity remain unresolved. Moreover, septins may sense membrane curvature and deform membranes, facilitating additional protein binding—a feedback process observed in synthetic systems (*12*, *13*). This membrane-deforming ability is likely due to the multivalent arrangement of AH domains, which serve as tether points for septin scaffold assembly. Interestingly, while native septins display curvature preference, an isolated AH domain lacks curvature sensitivity (*14*). However, tandem AH constructs can restore the function (*15*), pointing to the importance of multivalent interactions and flanking effects in membrane binding and peptide configuration.

Amphipathic helices are short α-helical motifs characterized by a spatial separation of hydrophobic and hydrophilic residues, enabling them to insert into lipid bilayers and act as sensors or inducers of curvature (*16*, *17*). This structural feature underlies their widespread role in membrane dynamics across many protein families. For example, in BAR proteins, amphipathic helices contribute to the organization of higher-order structures (*18*). Association with the membrane stabilizes the AH helical structure, primarily through interactions between hydrophobic residues and lipid tails (*19*). AH–membrane interactions are also influenced by membrane curvature, lipid composition, and the presence of defects—regions where lipid tail exposure facilitates peptide insertion. Membrane defects on convex surfaces enhance AH folding, promoting its helical structure and altering defect distribution (*20*, *21*). Membrane defects, defined as exposed hydrophobic lipid tails, play a crucial role in protein engagement. Additionally, studies indicate that lipid composition and membrane curvature influence defect formation, thereby regulating protein recruitment (*22*, *23*). Although AH domains in various proteins are known to facilitate membrane binding, the specific mechanisms of binding and the contribution of multiple AH domains to the higher-order assembly remain poorly understood. Building upon the findings of our previous study (*14*), which showed that the helicity and charge distribution of isolated septin AH domains tune their membrane-binding affinity, this study explores how multiple AH domains interact in and near the membrane interface.

To address the unresolved question of how AH domains contribute to cooperative membrane binding and septin assembly, we used molecular dynamics (MD) simulations to investigate the behavior of the extended AH domain from Cdc12 from the *S. cerevisiae* septin complex. We included upstream and downstream residues to more closely reflect the native context of the Cdc12 C-terminal region (Fig. 1A). Simulations were initiated from two configurations: with the peptide placed in the solution above the membrane (unbound; Fig. 1B) or shallowly inserted into the bilayer (bound; Fig. 1C). We examined both single-peptide and two-peptide systems to explore potential cooperative effects. We begin by characterizing the structural behavior of a single AH peptide that interacts with planar lipid bilayers. Then, we analyzed multi-peptide simulations to assess how interpeptide interactions may influence membrane association. The results suggest that the presence of a membrane-bound peptide can modulate the orientation and positioning of another peptide, likely through specific stabilizing interactions. Finally, we simulated covalently linked AH domains—referred to as tandem constructs (*15*) —to examine how connectivity influences structural stability. While palindromic CN-NC configurations exhibit instability, containing charged C-terminal domains stabilized the extended tandem structure. These extended tandem AHs may represent a simple model of septin with a minimal linker between two AH domains and C-terminals of Cdc12. Together, these simulations provide molecular-level insights into how AH domains may cooperatively engage membranes and offer a framework for understanding or engineering membrane-binding peptides with tunable properties.

**Figure 1.**
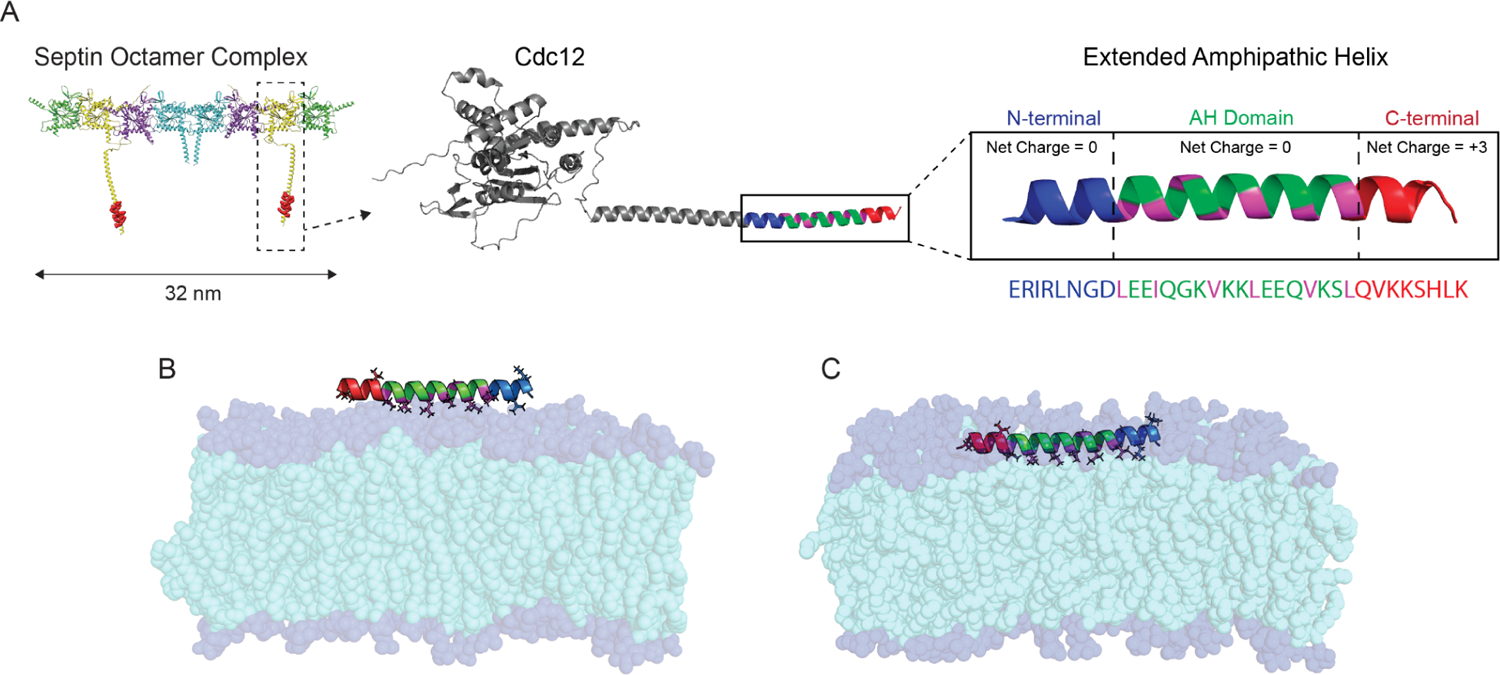
Septin structure and initial configurations of the extended AH with a lipid bilayer. (A) PDB structure of a nanoscale septin oligomer with a zoomed-in view of the Cdc12 subunit and its Amphipathic Helix (AH) domain with adjacent residues. The extended AH regions are color-coded: N-terminus (blue), AH domain (green), hydrophobic residues (magenta), and C-terminus (red). The corresponding amino acid sequence is also shown. (B) Initial configuration of the unbound state: the extended AH peptide is placed in solution above the lipid bilayer. (C) Initial configuration of the bound state: the extended AH is shallowly inserted into the membrane. The AH hydrophobic face is aligned with the lipid tails. Lipid hydrophobic tails are shown in cyan and phosphate headgroups in dark blue. Lipids are rendered semi-transparent to highlight the peptide orientation and residue types.

## MATERIALS AND METHODS

This computational study of a system with peptides and lipid bilayer is performed using GROMACS 2022.3 (*24*). The amino-acid sequences were derived from the septin protein in budding yeast S. cerevisiae Cdc12, which contains the amphipathic helix (AH) domain (*15*). We prepared the initial structure of the peptides using Pymol. Then the complete system, including the peptide, lipid bilayer, and the solution of water and ions, was assembled using the CHARMM-GUI web server (*25*) utilizing the CHARMM36m force field (*26*, *27*). The salt concentration in the solution is 50 mM KCl, and the temperature is 300 K. We set a timestep of 2 fs, with Van der Waals (VDW) interactions switched between 1.0 and 1.2 nm, and particle mesh Ewald was used at long distances to compute the electrostatic interactions (*28*). We used the LINCS algorithm to constrain hydrogen bond lengths (*29*). Evolution of the system was performed by the Nose-Hoover thermostat with a coupling constant (τ*t*) of 1 ps and Parrinello-Rahman barostat at a reference pressure of 1.0 bar and compressibility of 4.5×10^−5^ with a coupling constant of (τ*p*) of 5 ps. The pressure was coupled semi-isotropically in XY for membrane simulations and isotropically for peptide-only simulations.

### Single extended AH domain

The extended AHs, including the AH domain in the middle (18 residues) and extended residues on both ends (8 residues on each side) create a 34-residue peptide we call extended AH (*ERIRLNGDLEEIQGKVKKLEEQVKSLQVKKSHLK*). This sequence is located on the C-terminal of S. cerevisiae Cdc12 (Fig. 1). The net charge of the middle AH domain and eight upstream amino acids on the N-terminal are neutral, but the eight downstream amino acid sequence on the C-terminal has a +3 net charge under our experimental conditions (pH = 7.4). First, we build an idealized α helix of these 34 amino acids in PyMOL (*30*) and utilize CHARMM GUI to add neutral cappings for both N-terminal (NNUE) and C-terminals (CT2).

### Lipid bilayer

To build the lipid bilayer, we utilized the CHARMM-GUI membrane builder (*31*, *32*) (Jo et al., 2009; Wu et al., 2014). The generated bilayer was composed of a mixture of DOPC:PLPI lipids with a 75:25 proportion. The XY size of the simulation box is set to be roughly 11×11 nm^2^ and solvated with at least 5 nm of water (TIP3P forcefield) above and below the bilayer. The extended AHs are aligned with the bilayer in CHARMM-GUI so that the ILE12 and LEU19 residues are oriented to face downward to the interior, the hydrophobic portion of the bilayer (Fig. 1B and C). For the bound initial structure, we create a system of the lipid bilayer and place the extended AH parallel and at a z=15 Å distance to the membrane midline. For the unbound initial structure, the distance of the extended AH center of mass from the midline is z=30 Å. Systems were relaxed and equilibrated using the CHARMM-GUI steps (*32*).

### Two peptides

#### An unbound and a bound extended AH on the lipid bilayer

We create a system of the lipid bilayer with a bound extended AH and an extended AH unbound in the solution above the bilayer: to build this, we took the last frame of single-bound peptide simulations and added another extended AH to the solution. The unbound peptide is located parallel at the 30 Å distance from the bound peptide. Then, we deleted overlapping water and ion atoms within the distance of 7 Å of the unbound peptide atoms. In this method, the bound peptide is already in the bending shape with some hydrophobic residues on the extended parts facing outward of the bilayer.

#### Two bound extended AHs on the bilayer

The initial structure includes two bound extended AHs in the parallel conformation (C-terminal of both in one direction) with a 15 Å distance relative to each other. We placed these two parallel peptides in a box of lipid bilayer, water, and ions using CHARMM GUI. Again, we align ILE12 and LEU19 orientation so that the hydrophobic residues of the AH domain face the hydrophobic lipid tails and hydrophilic residues face the polar lipid phosphate headgroups—both peptides at the height of z=15 Å from the bilayer midline.

### Tandem AH domains

We connected two AH domains with the 10 amino acid sequence described in the experimental study (*15*). We generated three types of tandem structures:

1. **Tandem NC-NC AH domains:** linking the C-terminus of the first AH domain to the N-terminus of the second one. In this way, we get a 46-residue sequence called tandem AH NC-NC: *LEEIQGKVKKLEEQVKSLGSGSRSGSGSLEEIQGKVKKLEEQVKSL*.
2. **Tandem CN-NC AH domains:** linking the N-terminal of the first peptide to the N-terminal of the second AH domain. In other words, we flipped one of the AH domains to form a peptide with 46 amino acids that only the C-terminals of the AH domains are on the ends we call this structure tandem AH CN-NC or symmetric tandem that formed by palindromic arrangement of AH domains: *LSKVQEELKKVKGQIEELGSGSRSGSGSLEEIQGKVKKLEEQVKSL*.
3. **Tandem extended CN-NC AHs:** The palindromic arrangement of AHs plus the extended charged C-terminals on both ends. It forms the sequence of 62 amino acids: *KLHSKKVQLSKVQEELKKVKGQIEELGSGSRSGSGSLEEIQGKVKKLEEQVKSLQVKKSHLK*.

This design can be considered as a molecule that can mimic the septin protein structure with two AH domains and eight extended C-terminal residues on the two Cdc12s, regardless of the middle residue sequences and their length.

**Table 1.** Summary of molecular dynamics simulations.

| Simulation setup | Replicas,<br>duration (ns) | Peptide(s), cappings |
| --- | --- | --- |
| Single unbound | 4 × 400 | Extended AH, NNUE, CT2 |
| Single bound | 4 × 400 | Extended AH, NNUE, CT2 |
| Unbound and bound 30 Å distance | 8 × 400 | Extended AH, NNUE, CT2 |
| Both bound, 15 Å distance | 4 × 400 | Extended AH, NNUE, CT2 |
| Tandem NC-NC, unbound | 4 × 400 | Linked AH domains, NNUE, CT2 |
| Tandem CN-NC, unbound | 4 × 400 | Linked AH domains, CT2, CT2 |
| Tandem extended CN-NC, unbound | 4 × 400 | Linked Extended AH, CT2, CT2 |

#### Post-processing and analysis of the simulation outputs

A summary of molecular dynamics simulations can be found in Table 3.1. We analyzed the simulation trajectories using Python scripting and packages of MDAnalysis (*33*), MDtraj (*34*), and membrane-curvature (*35*). Visualizations were done by PyMOL and VMD (*36*).

Salt bridge interactions between peptide residues were analyzed using both VMD and PyMOL tools. VMD’s salt bridge plugin was used for trajectory-based analysis of inter-residue electrostatic contacts, while PyMOL’s salt bridge analysis was applied to representative structural snapshots for visualization and verification.

Amino acid sequences of the extended amphipathic helix (AH) domains from selected species were retrieved from the UniProt database(*37*). Multiple sequence alignment was performed using Jalview (*38*), and the aligned sequences were visualized with WebLogo (*39*) to highlight conserved positions. Charged residues were specifically examined to assess conservation patterns relevant to inter-peptide interactions and salt bridge formation.

To analyze the local bending of the peptide backbone, we used the Bendix plugin within the VMD package (*40*) to compute and visualize local bending angles —quantitative measures of how sharply the peptide backbone deviates from linearity at specific points along its contour, which are closely related to local curvature.. In this analysis, we set the side parameter in Bendix to 7.2 Å, reflecting the chord length measured between pairs of Cα atoms, along the helical backbone, to define each segment for curvature calculation. This corresponds approximately to the straight-line distance between Cα atoms separated by ∼3 to 4 residues in an α-helix, effectively covering a full helical turn. This setting balances resolution and smoothness in the curvature profile: smaller values increase local sensitivity but may amplify thermal noise, whereas larger values average out finer structural features. To quantify the overall curvature, we fitted a circle to the backbone alpha carbon positions of the peptide projected onto its best-fit plane. This plane was determined using principal component analysis (PCA).

For lipid organization analysis, we used the FatSlim package (*41*) to compute two key structural properties of the membrane: the area per lipid (APL), and the nematic order parameter of lipid tails. We generated maps of APL across the membrane plane in the form of two-dimensional heatmaps highlighting local variations in lipid packing density. These maps help visualize regions of tight or loose lipid organization that may be associated with protein insertion effects.

To assess the alignment of the lipid tails, we calculated the nematic order parameter, which quantifies the degree of orientational ordering of hydrocarbon tails relative to the membrane normal. For each lipid, the nematic order is defined as the 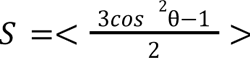, where *θ* is the angle between a carbon-carbon (C–C) bond vector along the lipid tail and the membrane normal vector. The angle brackets denote averaging over all bond vectors within a single lipid. This approach enables us to resolve local variations in membrane ordering by capturing differences in lipid tail orientation across the bilayer plane.

## RESULTS

### The extended AH peptide adopts a stable, curved conformation when bound to the membrane

We began by analyzing the behavior of a single membrane-bound extended AH peptide composed of 34 residues, including N- and C-terminal extensions flanking the core amphipathic helix (AH). Across all four simulation replicas, the bound peptide consistently adopted a stable, curved, smile-like conformation throughout the trajectory, as shown in a simulation snapshot in Figure 2A. This curvature appears to be a consequence of the peptide’s domain-specific interaction with the membrane. The central ∼18 residues correspond to the core amphipathic helix (AH), which exhibits a clear spatial segregation of hydrophobic and polar side chains, enabling it to stably associate with the membrane surface via hydrophobic insertion. In contrast, the N- and C-terminal extensions flanking the AH do not follow a regular amphipathic pattern and lack the structural features required for stable membrane insertion. As a result, the central AH region remains closely bound to the membrane surface, while the terminal segments are lack this pattern and tend to bend upward toward the aqueous phase. This asymmetric interaction along the peptide length generates a net bending moment, resulting in the observed smile-like conformation. Supporting this interpretation, simulations of the isolated AH segment alone (without the flanking regions) do not exhibit significant curvature, highlighting the importance of the full peptide architecture in driving this membrane-bound conformation.

**Figure 2.**
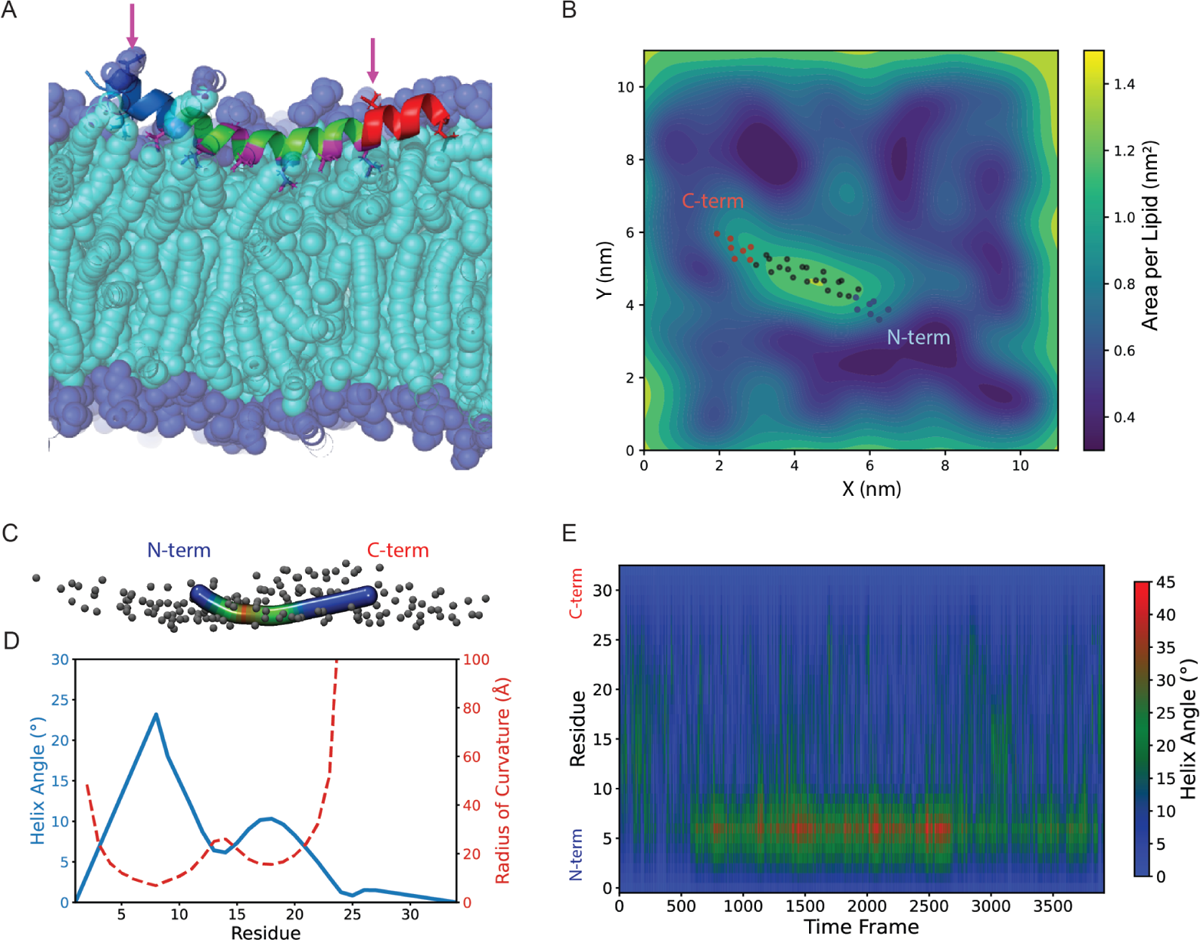
Bound peptide adopts a stable and smile-like curvature in the membrane. (A) MD simulation snapshot of a membrane-bound extended AH, with the N-terminus in blue and the C-terminus in red. Arrows indicate hydrophobic residues in the extended regions that face outward toward the solution, unlike the hydrophobic face of the AH region that orients inward. (B) Snapshot of the spatial map of area per lipid (APL) in the membrane upper leaflet. The presence of the peptide increases local APL near its binding site. The dots indicate the position of the backbone alpha carbons. (C) Peptide conformation color-coded by local bending angle using the Bendix tool. (D) Helix angle and calculated curvature corresponding to the Bendix snapshot. (E) The Helix angle map over simulation time shows stability in the local bending of the membrane-bound peptide. N-terminal residues bend more than those on the C-terminal side.

The presence of the bound peptide perturbs the surrounding lipid environment. The area per lipid (APL) map (Fig. 2B) shows local lipid packing defects near the binding site compared to distant unperturbed regions, indicating membrane remodeling induced by the bound peptide.

Unlike the amphipathic helix (AH) region, the extended residues do not maintain a clear spatial segregation of hydrophobic and hydrophilic side chains. As a result, not all hydrophobic residues on the extended domains can orient toward the membrane lipid tails. These hydrophobic residues that face outward are highlighted by arrows in Figure 2A. This contrasting amphipathic character between the AH and extended domains may contribute to the overall curvature of the bound peptide. The radius of the fitted circle, representing the overall radius of curvature, was found to be approximately 60 Å in all replicas (see Fig. S1). This bending was not observed in our previous simulations of the shorter AH domain comprising only 18 residues (Edelmaier et al., 2025), which remained largely straight when membrane-bound. The introduction of extended regions, which lack clear amphipathic character, gives rise to structural asymmetry that may contribute to the observed bending within the membrane environment.

Figure 2C shows a representative snapshot of the peptide, color-coded based on the local bending angle in the VMD Bendix tool. The corresponding local bending angle and radius of curvature of the snapshot is shown in Figure 2D. The peptide bending is not evenly distributed: the N-terminal region exhibits greater bending than the C-terminal region, and this asymmetric bending pattern remains stable throughout the simulations. This trend is visualized in the heatmap of the helix angle per residue over time in Figure 2E (also see Figure S2 for additional replicas). Notably, the highest degree of bending in the membrane-bound peptide occurs near residue G7, located at the interface between the N-terminal extension and the start of the AH domain. Since the AH domain spans residues 9 to 26, this sharp bend at the boundary likely reflects an interface effect, as further illustrated by the helix angle time average profile shown in Fig. S3.

### Inter-peptide interactions modulate the positioning and orientation of the unbound peptide

To test whether a membrane-bound peptide can facilitate the recruitment of additional peptides, we simulated a system containing two extended AH peptides: one initially bound to the membrane and the other initially placed in solution. As a control, we also analyzed a system of a single unbound peptide near the membrane. The Z-component of the center of mass (Z-COM) distribution for the unbound peptide shows a clear peak difference between the positioning of the unbound peptide in the single-(orange) and two-peptide (green) systems in Figure 3A. The gray curve, representing the distribution of phosphate headgroup positions, serves as a membrane reference. In the two-peptide system, the unbound peptide is more likely to reside closer to the membrane, as shown by a higher peak in the Z-COM histogram compared to the single-peptide system. This is further supported by Figure S4, which shows the AH domain Z-COM position over simulation time for each replica of the single-peptide system (*n* = 4) and the two-peptide system (*n* = 8). In the two-peptide systems, the unbound peptide remains closer to the membrane with reduced vertical fluctuations compared to the single-peptide systems that exhibit more variability in terms of membrane proximity.

**Figure 3.**
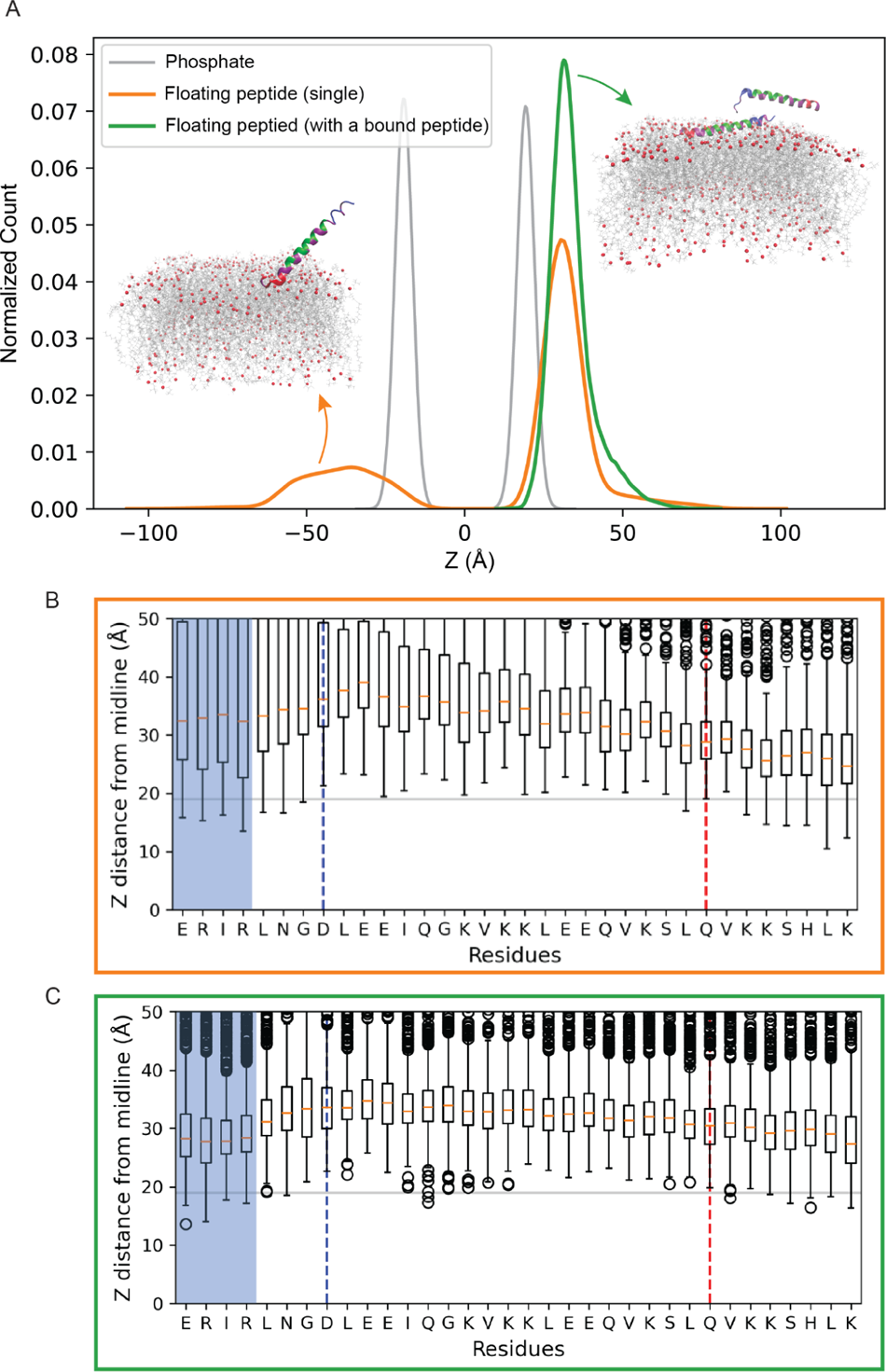
The presence of a membrane-bound peptide modulates the positioning and orientation of the unbound peptide toward the membrane. (A) Distribution of the Z-component of the center of mass (COM) for the unbound peptide in systems with single (orange, n=4) and two peptides (green, n=4). The gray curve shows the distribution of Z-positions of lipid phosphate headgroups. Left inset: MD simulation snapshot showing the single unbound peptide approaches the membrane via the C-terminus (orange). Right inset: MD simulation snapshot showing an unbound and a bound peptide. The N-terminal region of the unbound peptide can interact with the bound peptide (green). (B) Box plot of the Z-COM of residues in the system with a single unbound peptide (n=4). The blue and red dashed lines separate the N- and C-terminal regions, respectively. (C) Box plot of the Z-COM of residues of the unbound peptide in the multiple-peptide system. The first four N-terminal residues (blue box) show a closer approach to the membrane, consistent with interaction with the membrane-bound peptide.

To better understand the orientation of the unbound peptide toward the membrane, we examined the Z-COM positions of residues for the unbound peptide in both systems (Fig. 3B, C; see also Fig. S5). In the single-peptide system (Fig. 3B), the C-terminal region (red) approaches the membrane more closely than the N-terminal region (blue), likely driven by electrostatic attraction between the positively charged C-terminal residues and the negatively charged phosphate headgroups. This preferred approaching direction to the membrane via the C-terminus is also visible in the simulation snapshot (Fig. 3A, left inset).

On the other hand, in the two-peptide system (Fig. 3C), the N-terminal region—particularly the first four residues—shows closer proximity to the membrane than in the single-peptide case. When there is a bound peptide, both termini of the unbound peptide show proximity to the membrane. The N-terminal shift is likely mediated by direct interaction with the membrane-bound peptide, as shown in the snapshot (Fig. 3A, right inset). Notably, three of the first four N-terminal residues are charged (arginine and glutamic acid), suggesting a possible role for electrostatic interactions in mediating inter-peptide contacts and reorienting the unbound peptide.

### Charged residues form inter-peptide contacts in a preferred anti-parallel orientation

In the two-peptide simulations, although both termini of the unbound peptide occasionally come into proximity of the membrane surface, the N-terminus tends to remain closer to the bound peptide, whereas the C-terminus more often faces the membrane, engaging with phosphate headgroups. These observations raise the question of whether stable inter-peptide contacts form and whether specific conserved residues (see Figure 6B)—particularly charged ones—play a role in mediating these interactions.

To examine inter-peptide interactions, we computed contact maps based on distances between residues of the two peptides (Fig. 4A). We applied a 7.5 Å cutoff to the Cα distances to accommodate interactions mediated by long or flexible side chains. A zoomed-in view highlights a frequent contact between R4 of the floating peptide and E21 of the bound peptide. Figure 4B presents a domain-level contact map, coarse-grained across defined peptide regions and segmented into time windows for all replicas. The contact maps reveal that the N-terminal region of the floating peptide consistently tends to interact with the AH domain of the membrane-bound peptide. A closer look at the early time windows reveals that the initial dominant interaction occurs between the N-terminus of the bound peptide and the C-terminus of the unbound peptide, forming a preferred anti-parallel orientation. As the simulation progresses, the contact pattern stabilizes, with the most frequent interaction shifting to one between the N-terminus of the unbound peptide and the AH domain of the bound peptide. These interactions often involve charged residues in the N-terminus and within the AH domain, indicating a likely contribution of electrostatic interactions and potential salt bridge formation in stabilizing the anti-parallel alignment of the two peptides.

**Figure 4.**
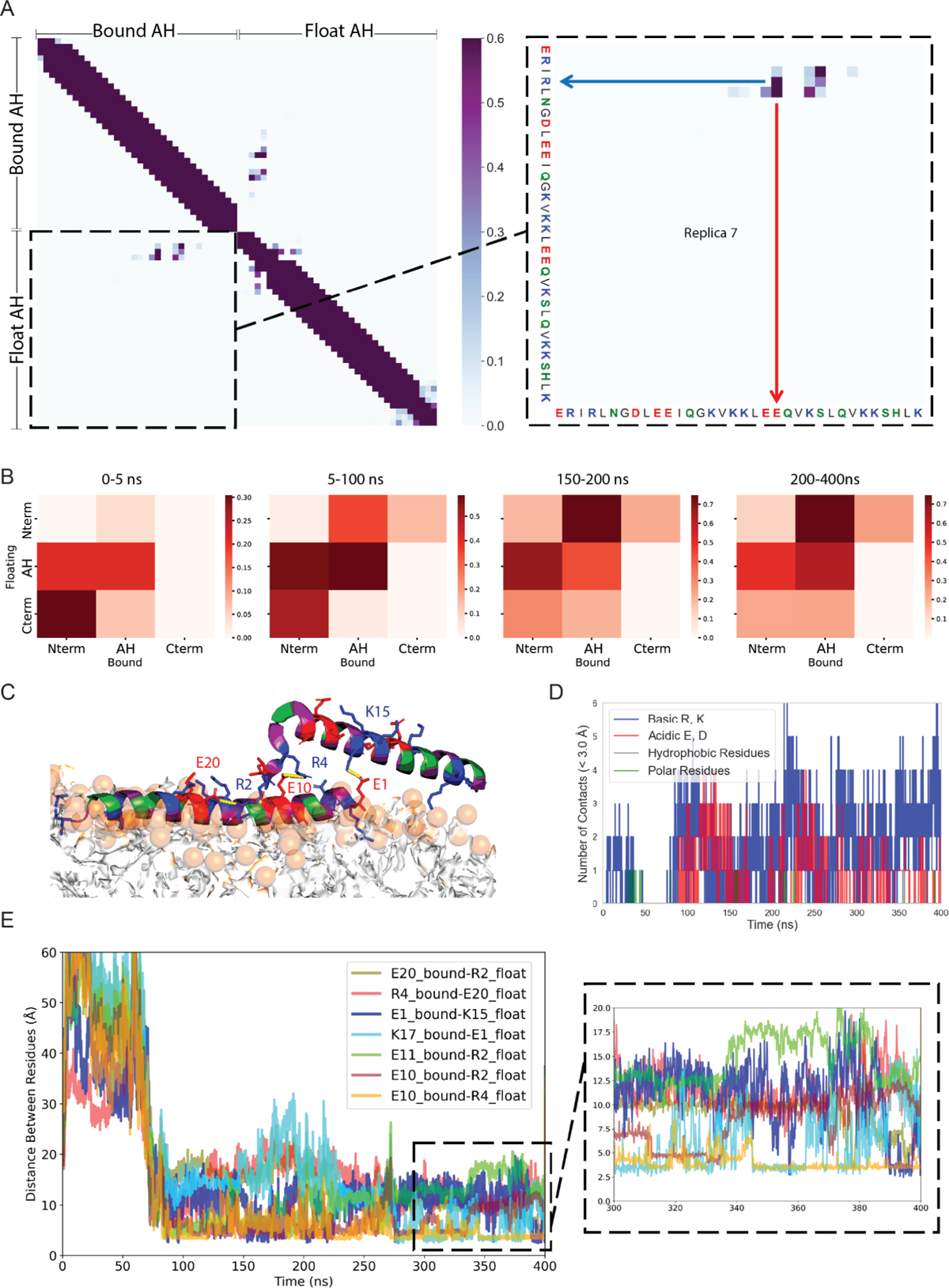
Salt bridge formation between charged amino acids promotes peptide-peptide interactions. (A) Contact map showing intra- and inter-peptide contacts. The zoomed-in inter-peptide region highlights a salt bridge between charged residues R4 and E21. The N-terminal of the floating peptide preferentially interacts with the ending region of the AH domain of the bound peptide, indicating an antiparallel peptide orientation. (B) The domain-based coarse-grained contact maps for all replicas in different time windows show two peptides interacting in an anti-parallel configuration that reaches a steady state. (C) Simulation snapshot showing multiple salt bridges formed between charged residues of the floating and bound peptide. Basic residues are shown in blue, acidic in red, polar in green, and hydrophobic in magenta. (D) number of inter-peptide contacts between residue pairs of the floating and bound peptides. The majority of contacts involve charged residues. (E) Time evolution of residue pair distances for the top-ranked salt bridge contacts, revealing how the two peptides interact (replica 2). R and E residues form a stable salt bridge throughout the simulation.

Salt bridges—electrostatic interactions between oppositely charged residues—are known to play a key role in stabilizing protein-protein interactions and overall structural integrity (*42*). As illustrated in the simulation snapshot (Fig. 4C), multiple salt bridges form between acidic and basic residues, primarily involving the N-terminal region of one peptide and the AH domain of the other.

To expand our analysis beyond charged residues, we tracked different types of inter-peptide contacts —distinguishing acidic, basic, polar, and hydrophobic residues— based on the distance between heavy atoms. We applied a 3 Å cutoff to capture close side-chain interactions at each time frame. As shown in Figure 4D for replica 2 (see Fig. S6 for all replicas), charged residues dominate over polar or hydrophobic residues in forming close inter-peptide contacts. To further characterize these charged interactions, we identified the top-ranked charged residue pairs forming persistent contacts (Fig. 4E for replica 2 and Fig. S7 for other replicas). These diagrams indicate that interactions between arginine (R) and glutamic acid (E) are recurrent throughout the simulation trajectories.

We also investigated the structural behavior of tandem AH peptides in which two AH domains were connected by a 10-residue linker introduced in an experimental study (*15*). In this experimental construct, the C-terminus of the first AH domain is linked to the N-terminus of the second AH domain, forming a structure we refer to as a tandem NC-NC or anti-parallel AH domains. We performed all-atom simulations of this 46-residue tandem construct initially unbound. Although the simulation started with a straight conformation, the peptide gradually bent at the linker region, resulting in an anti-parallel arrangement of the two AH domains (Fig. S8A). A positively charged arginine in the linker interacted with the membrane surface, anchoring the peptide near the bilayer without full insertion. In this anti-parallel conformation, the two AH domains remained folded, with their hydrophobic faces aligned and helical content preserved throughout the simulation (Fig. S8D). In contrast, when the first AH domain is flipped to form a CN–NC tandem configuration in which the peptides adopt a parallel arrangement, the structure is less stable: the helices unfold, and therefore, the hydrophobic and hydrophilic faces are no longer clearly maintained (Fig. S8B). The contact map and helical content diagram (Fig. S8E) show lower helicity and random contacts in the parallel CN–NC case.

To test whether structural stability could be restored by mimicking more native septin features, we extended both AH domains with eight additional C-terminal residues from Cdc12, creating an “extended parallel” tandem peptide. In this system, both C-termini contributed to membrane interactions. Unlike the shorter parallel tandem, this construct retained helical and amphipathic structures throughout the simulation (Fig. S8C, F), likely due to the stabilizing influence of the charged C-terminal extensions. These results support our previous findings (*14*) that the addition of flanking regions enhanced the helicity and stability of the structure. Tandem simulation results suggest that both anti-parallel geometry and the presence of extended, charged termini contribute to the structural integrity of tandem AH domains. Notably, the anti-parallel configuration observed in these tandem simulations reinforces the stable orientation of the anti-parallel alignment observed in our two-peptide simulations.

### The positioning of bound peptides in the membrane and the organization of the lipids depend on the number of bound peptides

We further explored peptide–peptide interactions by analyzing systems containing two peptides initially bound and shallowly inserted into the lipid bilayer in a parallel conformation that the previous section showed is less favorable. Across eight simulation replicas, we observed that in 75% of the cases, both peptides remained bound to the membrane while maintaining contact with each other (Fig. 5A). In the remaining 25%, one peptide partially dissociated from the bilayer and shifted toward the other peptide, forming interactions primarily through its N-terminal region (Fig. 5B). The snapshot’s side and top views demonstrate how one peptide reorients away from the membrane to associate with another peptide.

**Figure 5.**
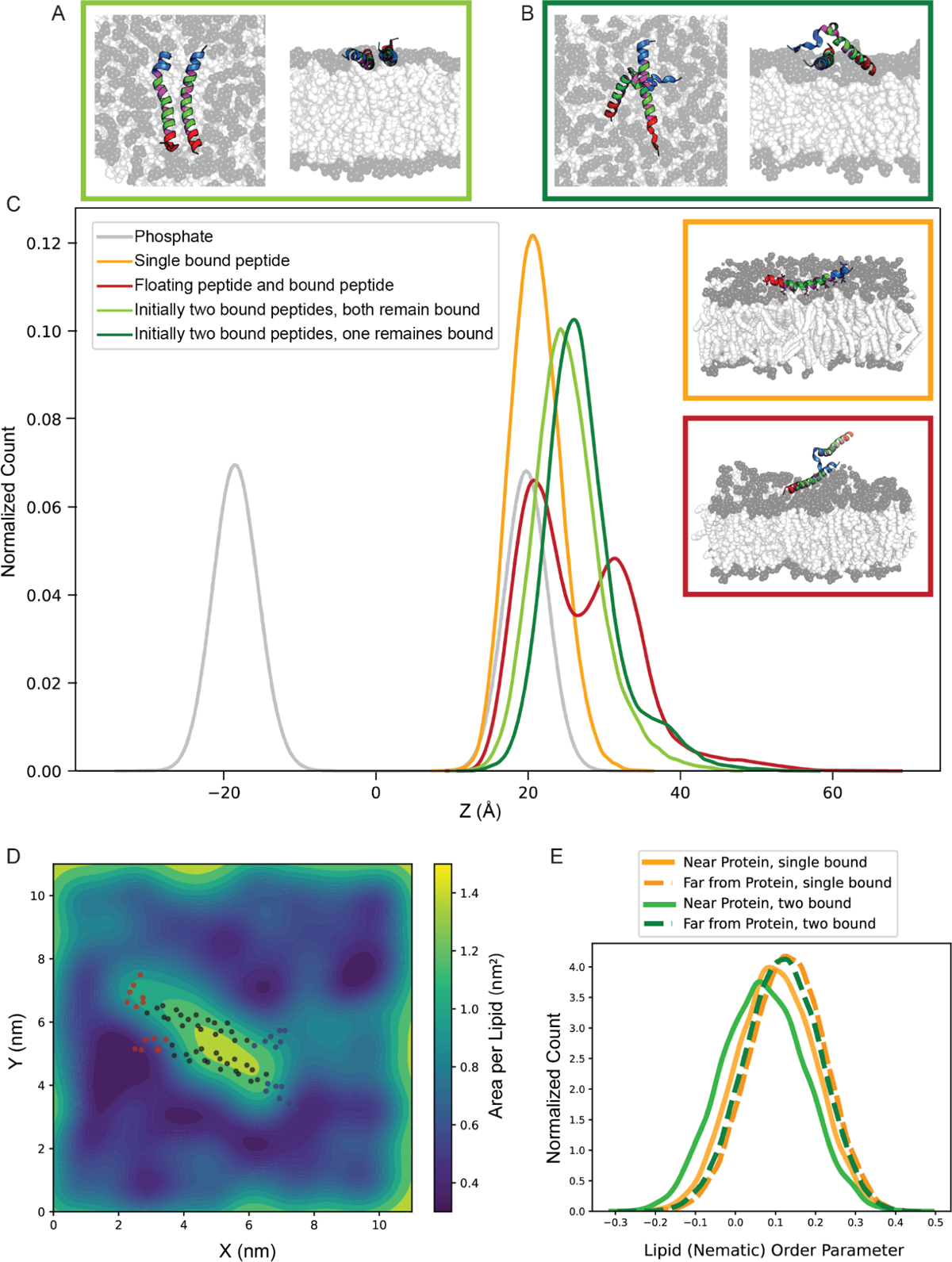
The position of bound peptides and the lipids arrangement change in single- and multiple-peptide systems. (A and B) Top and side view simulation snapshots of systems with the same initial structure of two parallel bound peptides show different behavior: (A) Both peptides remain bound in 400 ns simulation time (light green, 75% of replicas) (B) One of the peptides dissociates from the bilayer and interacts with the other peptide. (dark green, 25% of replicas). (C) Z-component of the center of mass distribution of the bound peptide in different systems. Inset: Simulation of snapshots of a single membrane-bound peptide (orange) and a system of bound and unbound peptides with lipid bilayer (red). (D) Spatial map of area per lipid (APL) in the membrane upper leaflet for two bound peptides initially placed in parallel (the non-favorable) orientation. The two peptides increase the local APL near the binding site and compact the distant lipids. The dots indicate the position of the backbone alpha carbons with N-terminal residues in blue and C-terminal residues in red. (E) Histogram of the lipids order parameter for systems with a single (orange) and two (green) bound peptides in the vicinity of (solid line) and far from (dashed line) the binding site.

In previous single-peptide simulations, the bound state appeared stable over the timescales sampled, with no evidence of spontaneous dissociation. In contrast, in 25% of two-peptide systems, the presence of a second peptide— even in the less favorable parallel orientation— altered this behavior, leading to the dissociation of one peptide from the membrane. This observation suggests that inter-peptide interactions can compete with or even override peptide–membrane associations. These outcomes imply that peptide–peptide contacts may provide a stabilizing influence that reshapes membrane binding dynamics in multi-peptide systems.

To provide an overview of peptide positioning relative to the membrane, we compared the Z-component of the center of mass distributions across different systems (Fig. 5C). In systems with two bound peptides (green), the Z-COM peak shifts to the right relative to that of the single-peptide system (orange), indicating that two bound peptides tend to reside slightly upward in the membrane. This upward shift becomes more pronounced over time, as shown in the time-resolved Z-COM distributions in Figure S9, suggesting the gradual displacement of the peptides during the simulation.

One possible explanation for this shift is lipid remodeling induced by increased peptide occupancy. The area per lipid (APL) map (Fig. 5D) reveals local lipid packing defects: APL values are lower near the peptides, while more distant regions become increasingly compacted. Compared to the more uniform lipid distribution in the single-peptide system (Fig. 2C), the two-peptide system displays more pronounced spatial heterogeneity, with more dark blue regions reflecting increased lipid density far from the binding site. This localized crowding, combined with inter-peptide interactions, may generate lateral pressure that slightly displaces the peptides from their original membrane-embedded positions. Consistent with this, the lipid order parameter distribution (Fig. 5E) shows that lipid tails become increasingly disordered as the number of bound peptides increases, particularly near the binding sites. The reduction in the order parameter likely reflects local perturbations of membrane structure caused by multiple interacting peptides. Together, these results suggest that increasing peptide occupancy alters both membrane organization and peptide positioning. In this context, a more curved membrane—by increasing available lipid area and reducing packing constraints—may better accommodate multiple bound peptides.

## DISCUSSIONS AND CONCLUSION

Our simulations suggest a mechanistic model for how septin AH domains interact to form assemblies on curved membranes. As illustrated in Figure 6A, a membrane-bound septin may facilitate the recruitment and alignment of neighboring septin complexes through salt bridge interactions between oppositely charged residues on their amphipathic helices. These interactions stabilize a preferred anti-parallel arrangement between AH domains, allowing for cooperative membrane association while preserving peptide helicity and amphipathic character. This configuration may represent an early step in septin filament formation, where lateral peptide–peptide contacts, stabilized by electrostatics, promote filament growth along the membrane surface. Curved membranes—by increasing lipid spacing and reducing local packing density—may further support such cooperative assembly by accommodating more AH domains at the interface.

**Figure 6.**
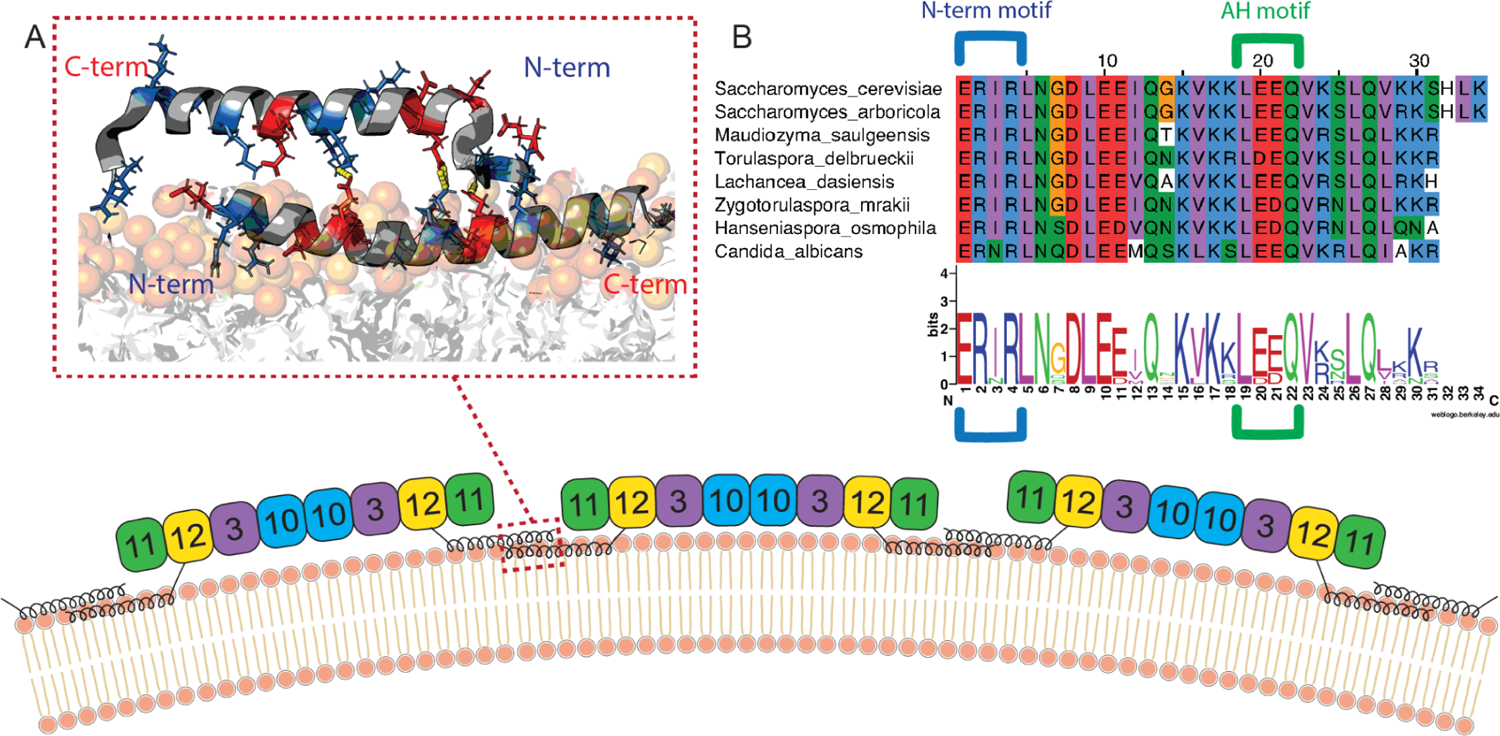
Proposed mechanism of septin polymerization on the curved lipid bilayer. (A) The bound septin facilitates the binding of neighboring septins through salt bridge interactions between their AH domains in anti-parallel, forming septin assemblies. (B) Sequence analysis shows conserved domains across different species, particularly charged residues that are detrimental to forming salt bridges and peptide-peptide contacts.

Importantly, the charged residues involved in these stabilizing interactions are evolutionarily conserved across septin homologs. Figure 6B presents a sequence alignment across species, emphasizing the conservation of key charged residues. Glutamic acid residues involved in salt bridge formation is located within the conserved LEEQV motif of the Cdc12 AH domain, while the interacting arginine in the N-terminal extension ERIR motif is also conserved across species (*15*, *37*, *43*). These conserved features may point to a universal strategy used by septins to stabilize their membrane-bound assemblies. Experimental validation through targeted mutagenesis of these residues would offer a valuable test of their role in inter-peptide interaction and filament formation.

Simulations of a single membrane-bound extended AH peptide revealed a stable smile-like curved conformation. This bending behavior is not observed in shorter constructs (*14*). This curvature is driven by the inclusion of non-amphipathic flanking regions that introduce asymmetry and enhance bending flexibility. Local curvature analysis revealed that the bending is asymmetric and most pronounced near the N-terminal interface. When a second peptide was added to the solution, its N-terminal region consistently approached the membrane-bound partner and formed inter-peptide interactions. In particular, salt bridges are frequently formed between arginines in the N-terminal extension of the floating peptide and glutamic acids within the AH domain of the membrane-bound peptide. These interactions potentially stabilize an anti-parallel configuration between the two peptides and suggest a mechanism of cooperative membrane association, where one bound peptide facilitates the recruitment and stabilization of another, reorienting and stabilizing it at the membrane interface.

Tandem AH simulations provided additional insights: peptides with NC–NC linkage adopted a folded, anti-parallel configuration stabilized by inter-domain contacts, while CN–NC tandem constructs showed unfolding and reduced stability. Interestingly, extending AH domains with C-terminal residues derived from Cdc12 restored stability even in parallel arrangements. This result highlights the role of terminal extensions in preserving helicity and amphipathic structure.

We also observed that increasing the number of bound peptides leads to structural changes in both peptide positioning and membrane organization. Two bound peptides induced local packing defects and lipid tail disorder, creating spatial heterogeneity in the membrane that differs from the single-peptide case. The area per lipid maps and lipid order parameters revealed lipid compaction at distal regions and increased disorder near the binding sites, which may generate lateral pressure that subtly displaces peptides from the membrane interior. This suggests that cooperative peptide binding may not only rely on inter-peptide contacts but also emerge from lipid-mediated effects that reorganize the membrane to accommodate multiple peptides. A curved membrane may better accommodate multiple bound peptides due to lipid spacing.

Looking forward, incorporating coarse-grained simulations or enhanced sampling techniques will be essential to extend accessible timescales and enable the observation of spontaneous binding events, thereby providing a more comprehensive understanding of septin-membrane association dynamics. Looking ahead, future work should explore multivalent peptide arrangements, curved membrane geometries, and longer simulation timescales to capture dynamic recruitment and filament formation more directly. Such studies may also inform the rational design of synthetic peptides and materials capable of mimicking biological curvature sensing and cooperative assembly mechanisms. In summary, our findings suggest that the presence of a membrane-bound peptide enhances the membrane proximity of the unbound peptide, possibly by creating a local interaction site or perturbing the lipid environment to favor association. These findings highlight the importance of both sequence features and membrane context in shaping AH domain interactions and suggest a plausible molecular mechanism by which septins assemble into filaments on the lipid bilayer.

## ACKNOWLEDGEMENTS

We would like to thank Pilar Cossio, Brandy N. Curtis, and Ellysa J. D. Vogt for their helpful discussions. We acknowledge support from the Alfred P. Sloan Foundation Matter-to-Life (grant no. G-2021-14197). R.F. acknowledges financial support in the form of a Cottrell Scholar Award (CS-CSA-2023-033), sponsored by Research Corporation for Science Advancement. The Flatiron Institute is a division of the Simons Foundation.

## AUTHOR CONTRIBUTIONS

S.M.M. performed simulations of systems with single, multiple, and tandem peptides, as well as results analysis and writing. A.S. helped with scripting and analysis. C.J.E. performed simulations of single AH domains and analysis. A.G., M.G.F., R.F., and S.J.K. provided helpful input and discussion. S.M.H. and E.N. supervised the work.

## DATA AND MATERIAL AVAILABILITY

All data supporting the findings of this study are available in the main text or the supplementary materials. Custom analysis code is publicly available at https://github.com/flatironinstitute/septin_AH_analysis. Representative simulation frames and processed data used to generate figures are included in the supplementary materials. Full simulation trajectories are available from the corresponding author upon reasonable request due to storage and bandwidth limitations.

