## Supplementary Information for "Study of Protein-Protein Interactions in Septin Assembly: Multiple amphipathic helix domains cooperate in binding to the lipid membrane"

**SUPPORTING MATERIAL**

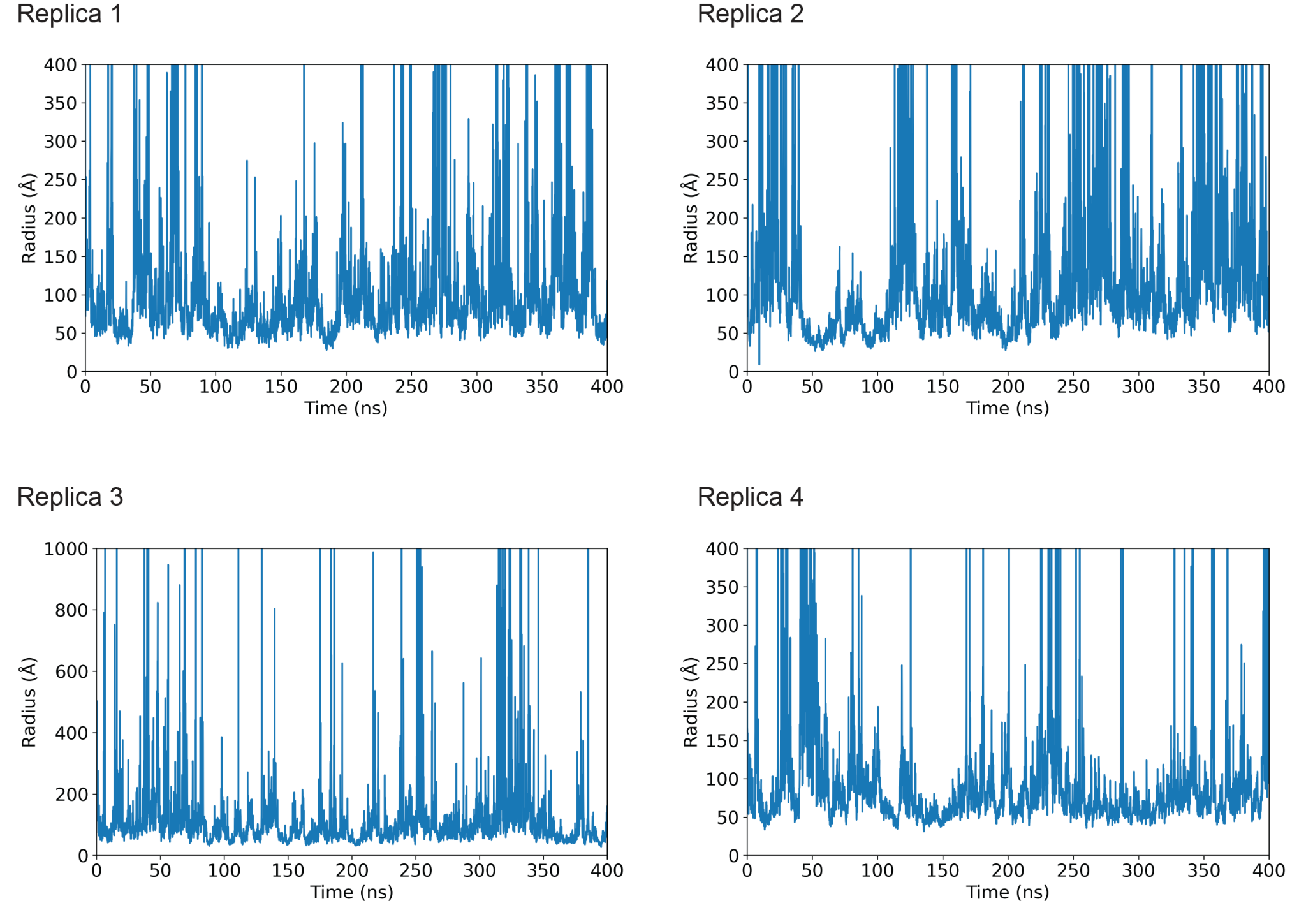
**Figure S1.** The overall radius of curvature of the extended bound peptide for different replicas. The peptide keeps ~60 Å curvature over the simulations.

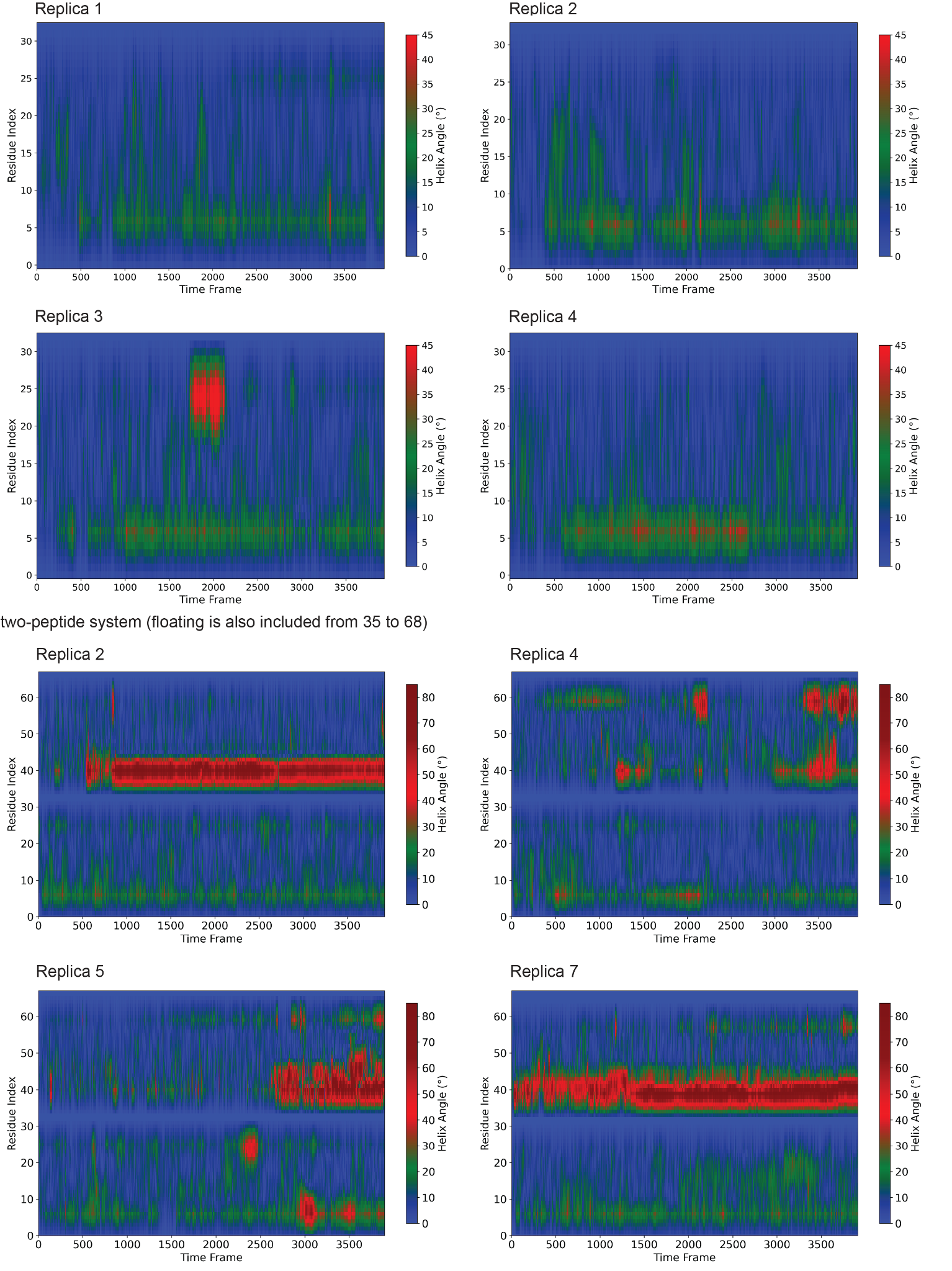

**Figure S2.** Bending angle heat map for single bound peptide in single and multiple systems for all replicas.

We repeated the Bendix analysis for this two-peptide system (Fig S2). The floating peptide—initially in solution—displays greater conformational flexibility, with the helix angle map showing more pronounced bending at the N-terminal residues compared to the C-terminal end. Comparing the average bending profiles of the membrane-bound peptide between single- and two-peptide systems (Fig. S3) reveals no considerable differences aside from the slight increase in C-terminal bending. These observations suggest that the primary driver of peptide curvature is the interaction with the membrane itself, particularly the distribution of hydrophobic and hydrophilic residues along the AH and flanking regions rather than inter-peptide contacts.

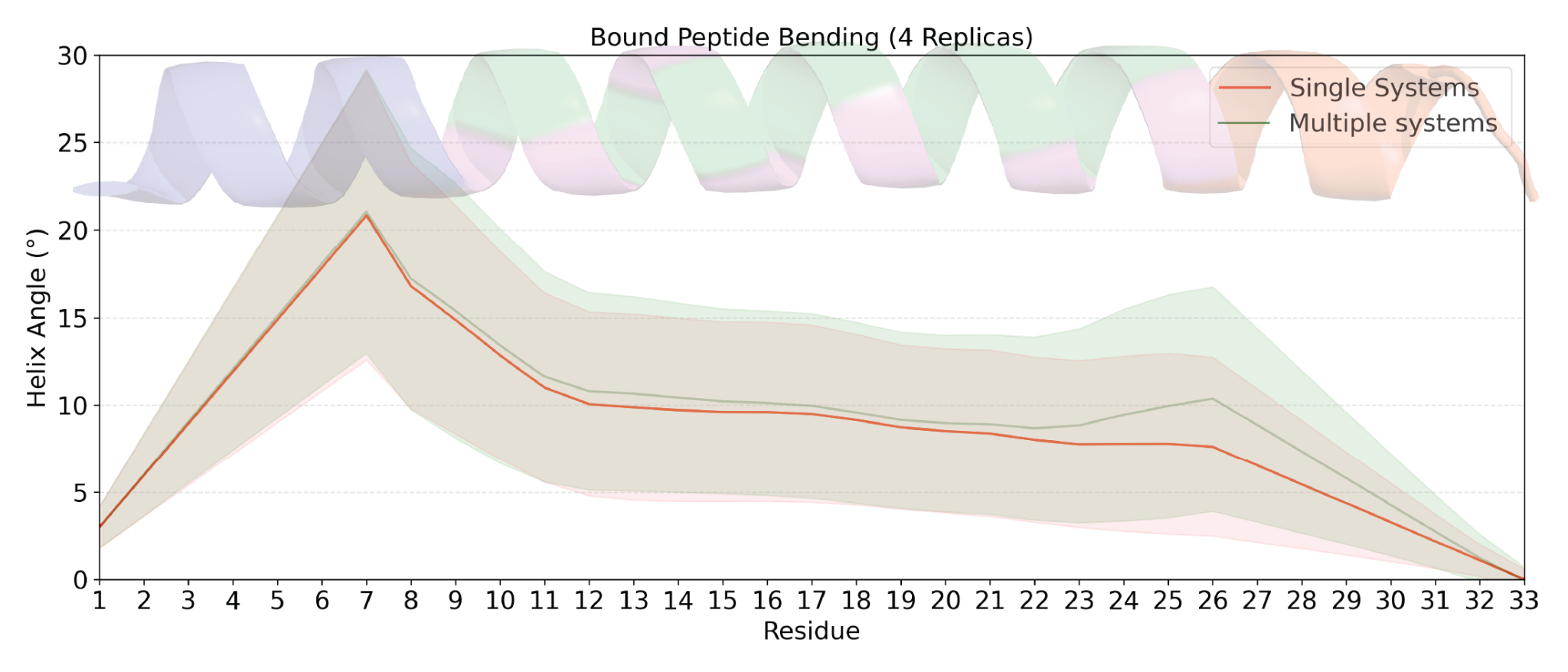
**Figure S3.** Average bending angle profile of the bound peptide over all four replicas in single- and two-peptide systems.

The highest bending is observed at the interface between the AH domain and N-terminal extention. To complement the local analysis, we also computed the global curvature of the peptide by fitting a best-fit circle to its smoothed backbone, yielding an overall radius of curvature of approximately 60 Å (Figure S1).

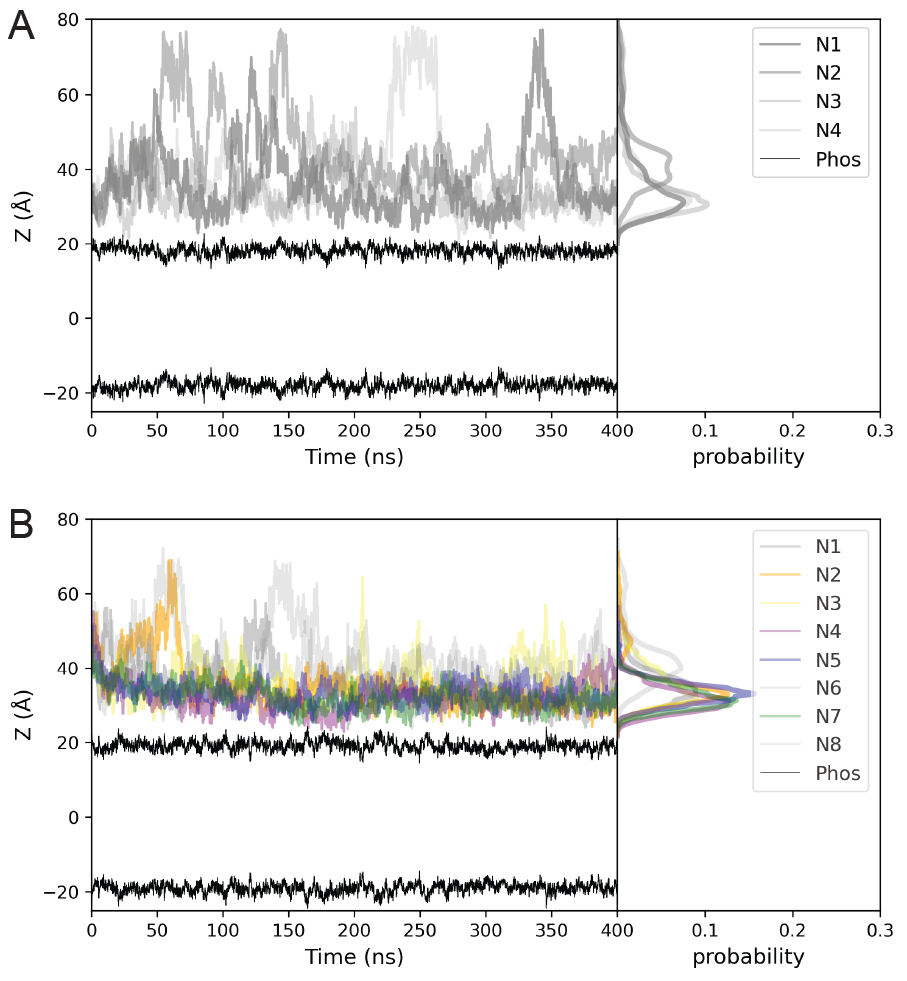

**Figure S4.** (A) Z-component of the center of mass of a single unbound peptide’s AH domain. The histogram is shown on the side for each replica (n=4). (B) Z-component of the center of mass of the unbound peptide’s AH domain in the presence of a bound peptide (n = 8). The histogram is shown on the side. Replicas with no interaction are colored in gray.

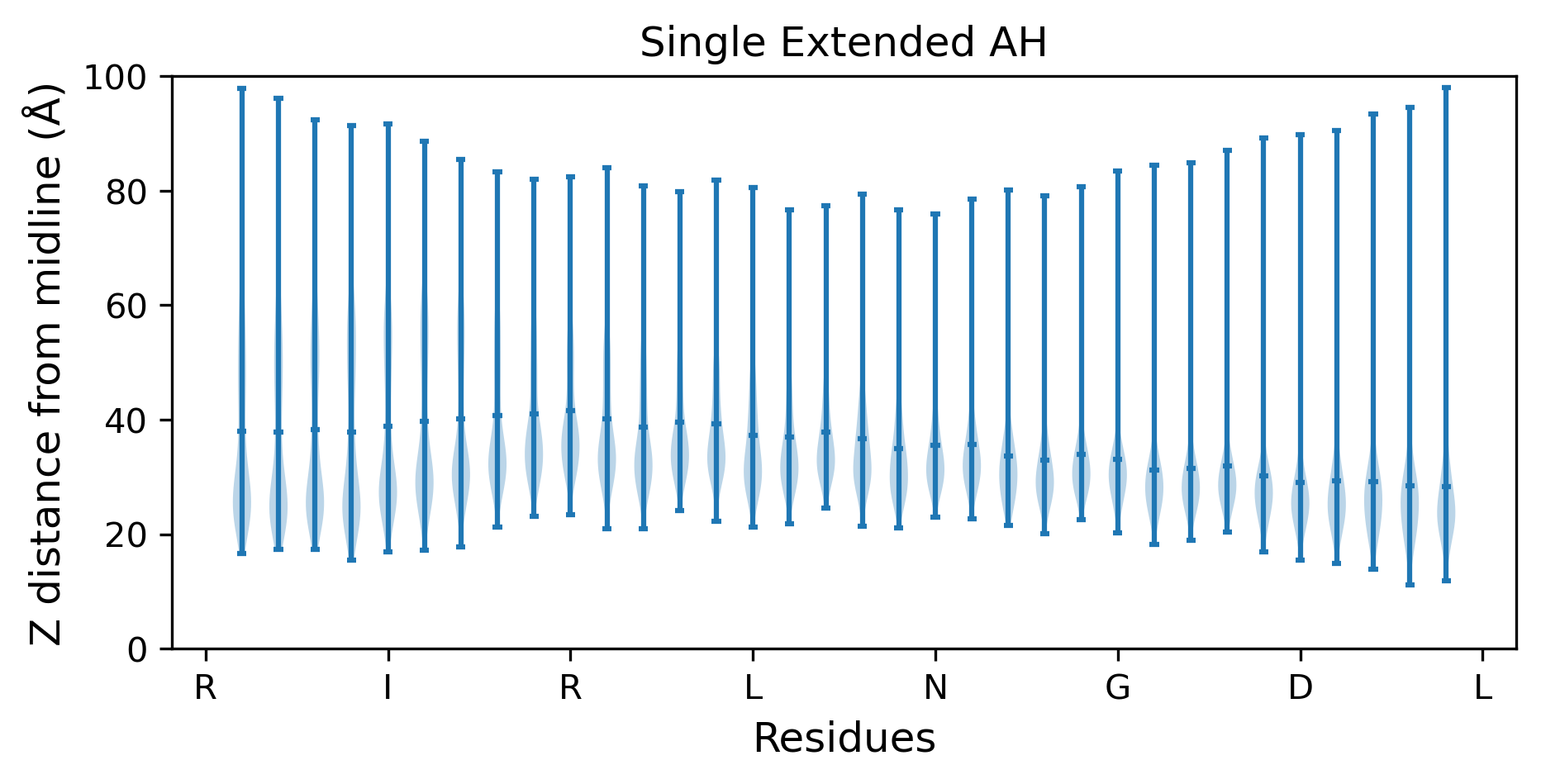

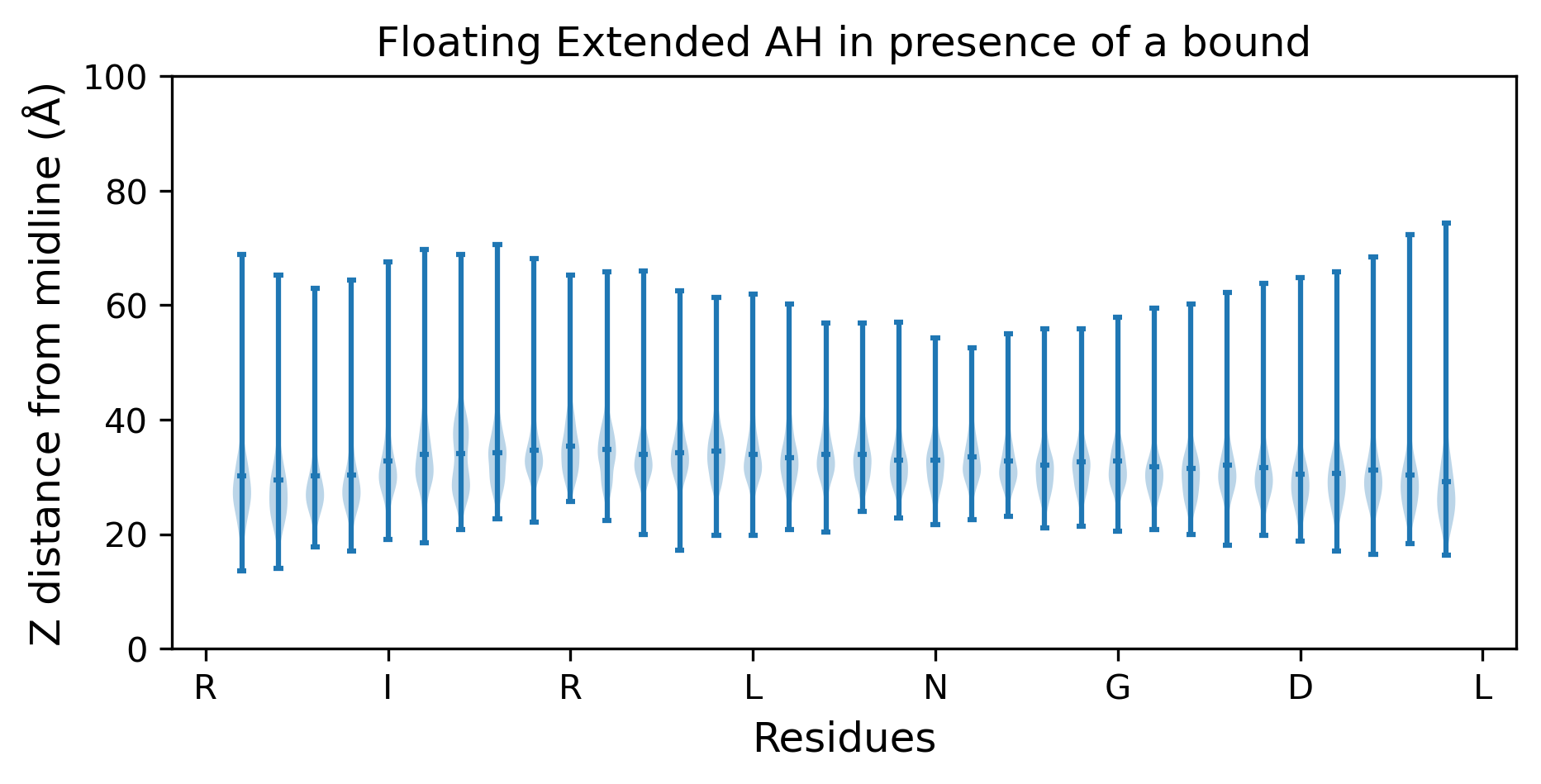

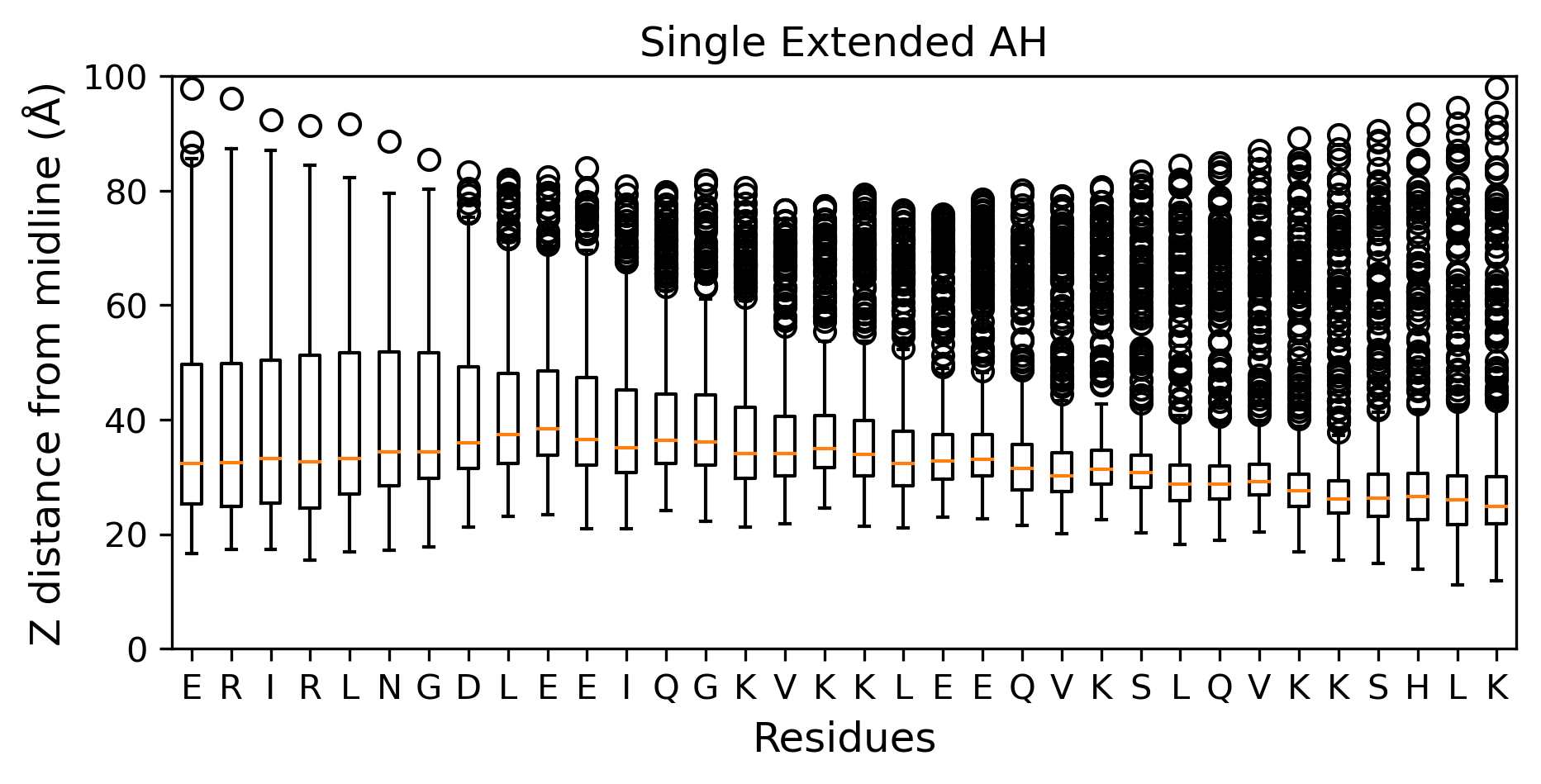

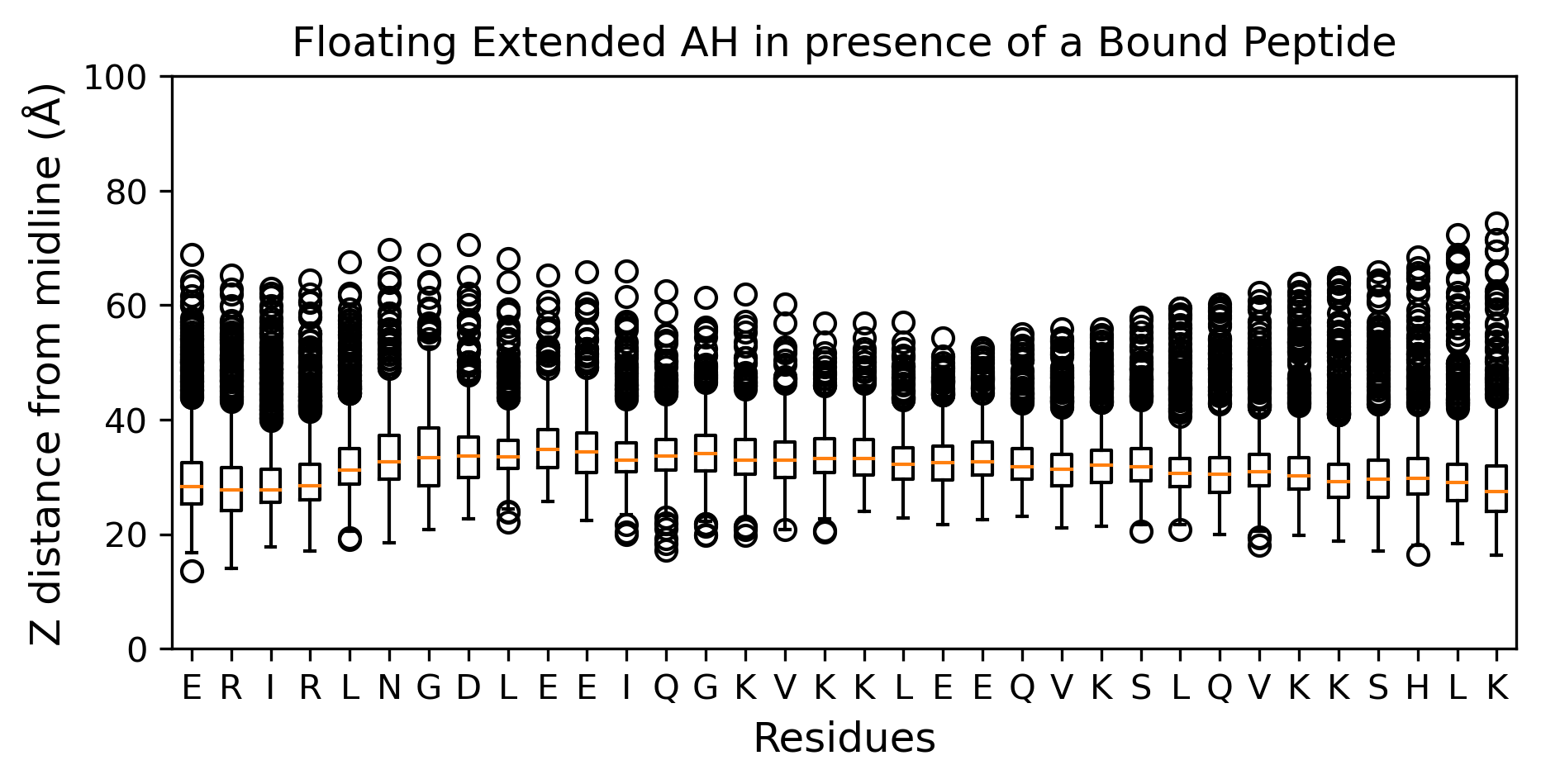

(A)	(B)

**Figure S5.** Violin and boxplot showing the median and deviation of the Z-component distance of the center of mass of the peptides from the membrane midline. (A) single unbound extended AH. (B) unbound extended AH in the presence of a bound peptide.

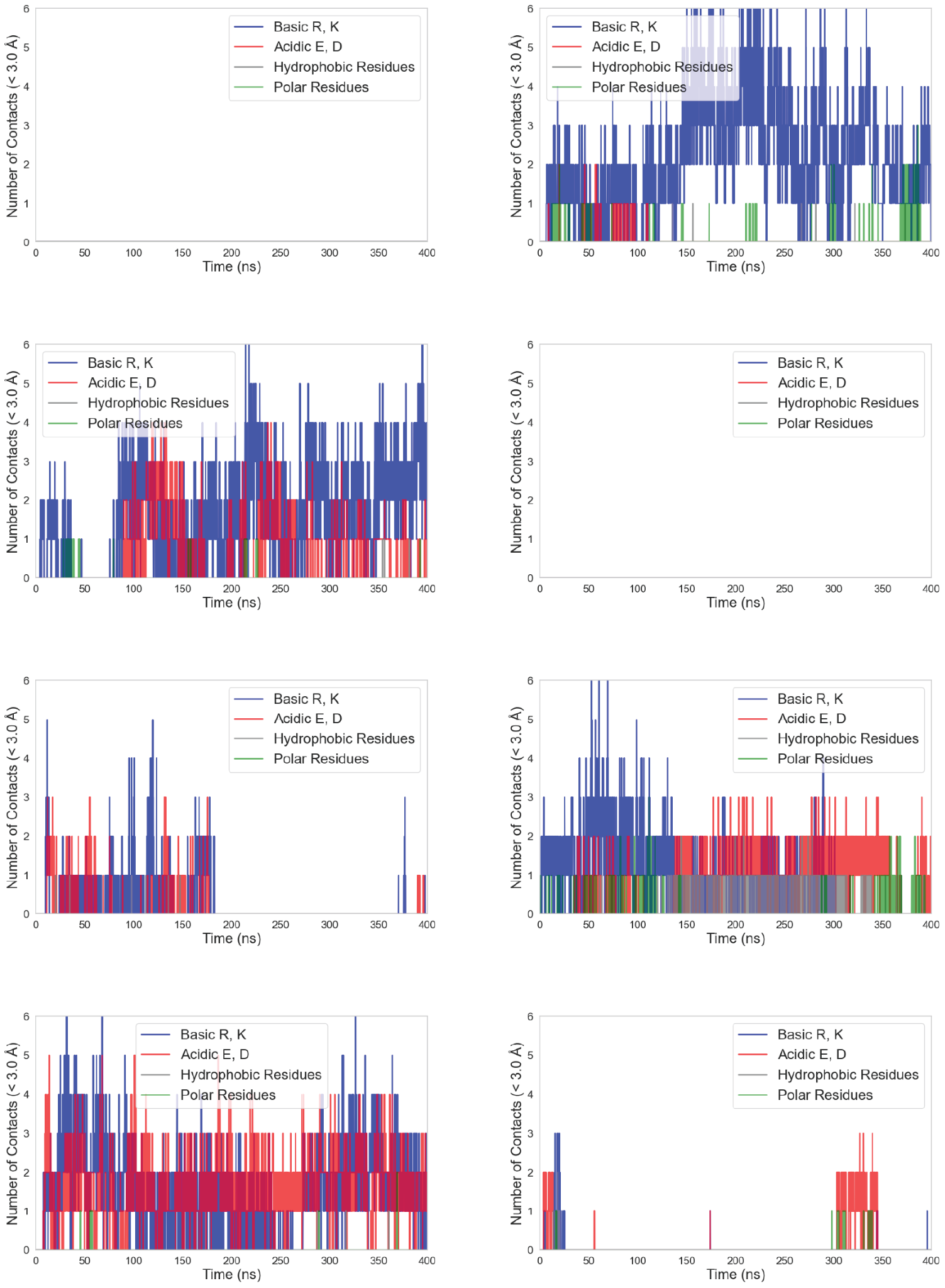

**Figure S6. number of contacts during simulation for each 8 replicas.** In replicas 1, 3, 6, 8 simulations, peptides could not find or interact with each other. For interacting peptides (replicas 2, 4, 5, 7), figures show how charged residues are dominant in forming contacts.

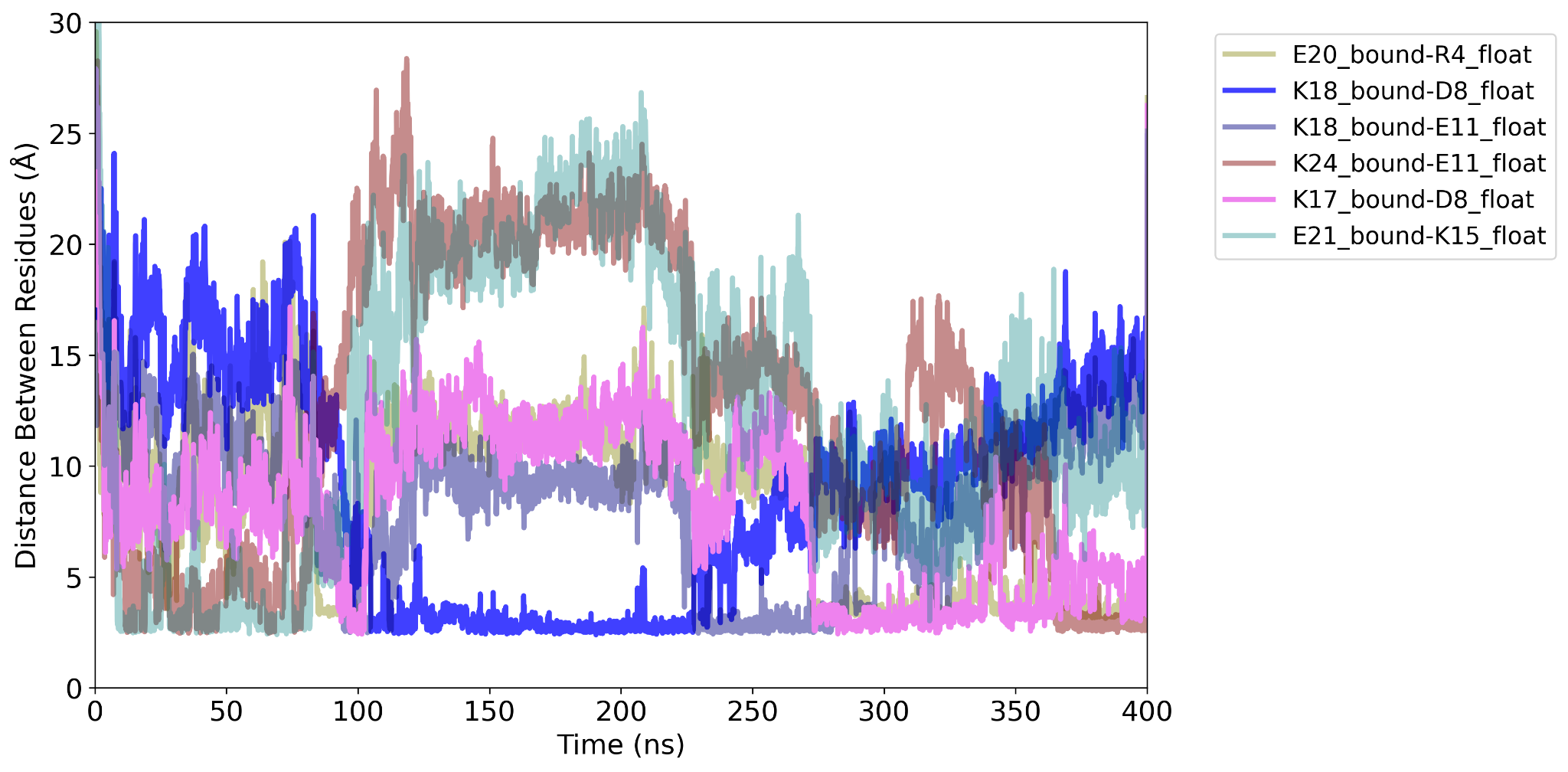

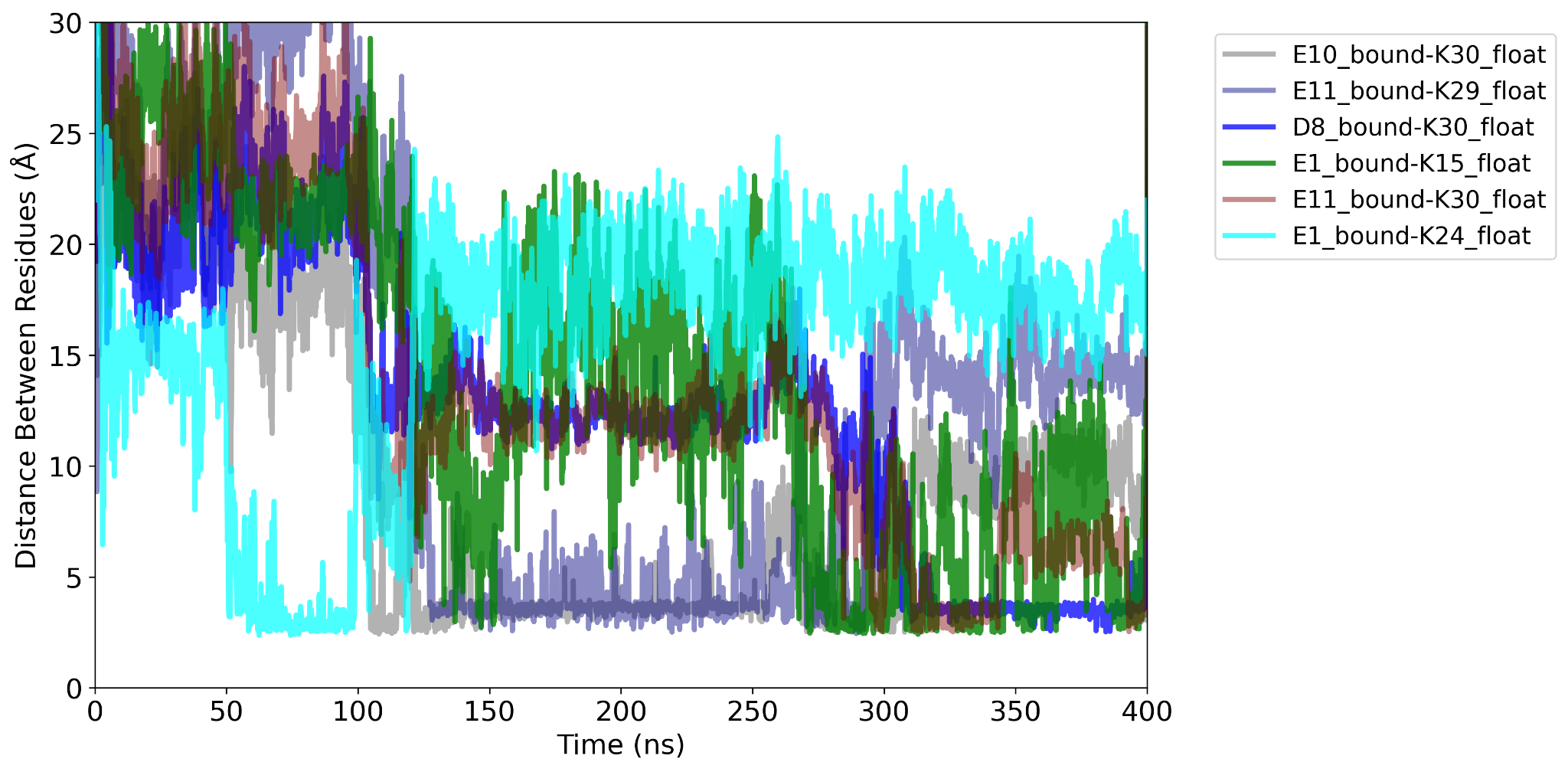

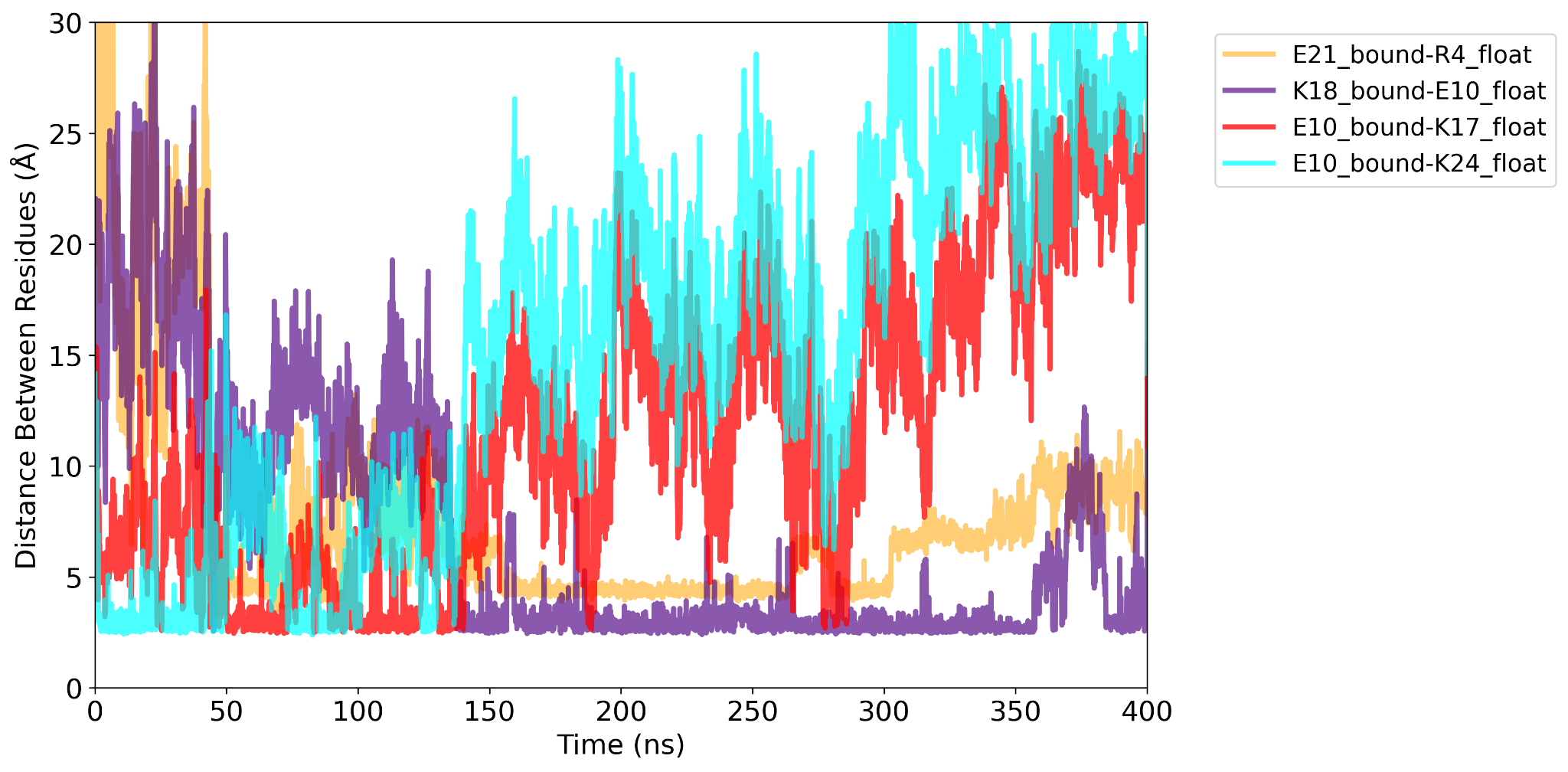

**Figure S7.** Time evolution of residue pair distances for the top-ranked salt bridge contacts, revealing how the two peptides interact (replicas 4, 5, and 7). R and E residues form a stable salt bridge throughout the simulation.

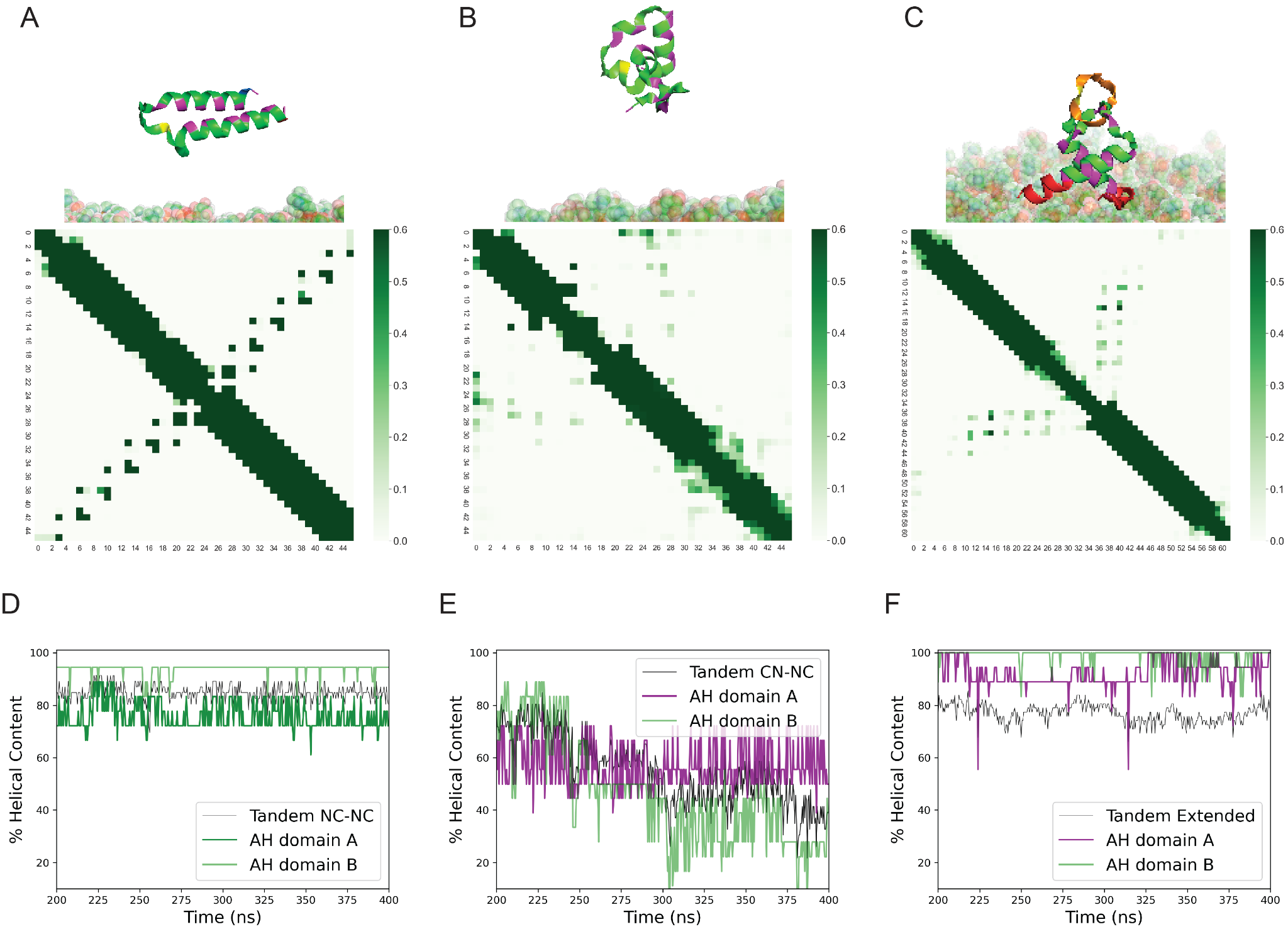

**Figure S8. The tandem peptide remains stable when AH domains are arranged in an antiparallel configuration or extended with charged C-terminals.** (A) Simulation snapshots illustrate how tandem AH domains in an NC-NC arrangement hinge at the central linker, forming a stable antiparallel configuration. Sharp features in the contact map indicate strong, stable interactions between anti-parallel AH domains. (B) Tandem AH domains in a CN-NC (parallel) configuration exhibit fluctuating interactions, with the contact map showing an unstable state. (C) Extended Tandem AHs with added charged C-terminals stabilize the parallel AHs in CN-NC configuration. (D-F) Helical content analysis reveals that stable states correspond to higher helical configurations. The NC configuration is shown in green, and the CN configuration is shown in purple for (D) anti-parallel, (E) parallel, and (F) extended parallel tandem AHs.

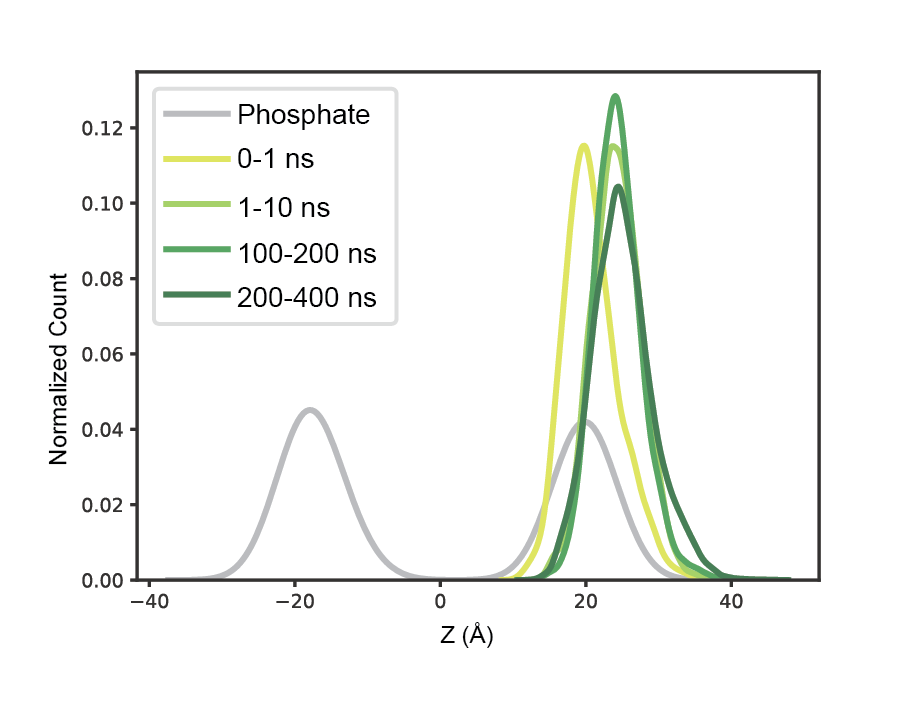

**Figure S9. Z-component of the center of mass distribution for peptides in simulations containing two membrane-bound peptides (*n* = 8), shown across different time windows.** As the simulation progresses, both peptides exhibit a slight upward shift, indicating gradual displacement away from the membrane interior
